# Exploring Pathogenic Mutation in Allosteric Proteins: the Prediction and Beyond

**DOI:** 10.1101/2024.03.23.586438

**Authors:** Huiling Zhang, Zhen Ju, Jingjing Zhang, Xijian Li, Hanyang Xiao, Xiaochuan Chen, Yuetong li, Xinran Wang, Yanjie Wei

## Abstract

Allosteric regulation that triggers the functional activity of a protein through conformational changes is an inherent function of the protein in numerous physiological and pathological scenarios. In the post-genomic era, a central challenge for disease genomes is the identification of the biological effects of specific somatic variants on allosteric proteins and the phenotypes they influence during the initiation and progression of diseases. Here, we analyzed more than 38539 mutations observed in 90 human genes with 740 allosteric protein chains. We found that existing allosteric protein mutations are associated with many diseases, but the clinical significance of the majority of mutations in allosteric proteins remains unclear. Next, we developed a machine-learning-based model for pathogenic mutation prediction of allosteric proteins based on the intrinsic characteristics of proteins and the prediction results from existed methods. When tested on the benchmark allosteric protein dataset, the proposed method achieves AUCs of 0.868 and AUPR of 0.894 on allosteric proteins. Furthermore, we explored the performance of existing methods in predicting the pathogenicity of mutations at allosteric sites and identified potential significant pathogenic mutations at allosteric sites using the proposed method. In summary, these findings illuminate the significance of allosteric mutation in disease processes, and contribute a valuable tool for the identification of pathogenic mutations as well as previously unknown disease-causing allosteric-protein-encoded genes.

## 1 Introduction

Allostery is a long-range effect with spatiotemporal features that exists in proteins. It describes a phenomenon where a specific site mutation or ligand binding triggers structural and/or dynamic changes in the protein at other sites[1]. The active site of the protein, which usually refers to the binding site for the natural substrate of enzymes or endogenous ligands of receptors, is called the orthosteric site. Since the specific site that induces allostery is different from the orthosteric site in space, it is called the allosteric site[2]. Gunasekaran et al.[3] predicted that allosteric regulation is an inherent attribute of the majority of proteins, except for rigid fibrous proteins. For proteins, this attribute means that the allosteric site can synergize with a distant region (such as the orthosteric site) in terms of motion or energy and transmit signals within the protein (i.e., allosteric pathways). Typically, the propagation of allosteric signals from the allosteric site to the functional orthosteric site is initiated by perturbations at the allosteric site, including the binding of modulators (ions, small molecules, proteins, DNA, and RNA)[4], post-translational modifications[5], and missense mutations[6]. Allosteric regulation affects the functionality of biomacromolecules in physiological conditions through allosteric sites, achieving precise regulation of numerous biological processes, including enzyme catalysis[7], gene expression[8], cellular differentiation[9], metabolism[10, 11], and organismal homeostasis[12]. Due to the ubiquity of allosteric control governing organismal life processes, allosteric regulation is considered as the “second secret” of life [13] and an essential approach to understanding cellular physiology and pathology[14]. Allosteric sites exhibit lower conservation compared to orthosteric sites[15], making them more susceptible to mutations. Mutations in allosteric sites (or allosteric signaling pathways) are closely associated with the occurrence and progression of many diseases[16, 17], and can also lead to resistance to allosteric drugs[18]. However, not all mutations in allosteric proteins are driver mutations in disease development; many mutations result in minor changes in protein function or manifest clinically as benign. Due to the uncertainty of their functional impact, most rare missense mutations in clinical genetic testing are classified as “uncertain” or imprecisely categorized in terms of the pathogenicity of a minority of mutations, leading to ambiguous clinical diagnosis in precision medicine, overtreatment, or missed clinical intervention opportunities. Accurate identification of pathogenic mutations in allosteric proteins not only provides effective clinical markers for precise diagnosis but also offers theoretical guidance for understanding disease pathogenesis, allosteric drug design, and personalized precision medicine, with significant medical value and social significance.

Currently, variants are commonly evaluated using the American College of Medical Genetics and Genomics (ACMG) criteria [19] or modifications thereof. According to the ACMG criteria, variants are categorized into five pathogenicity classes: “benign”, “likely benign”, “variant of uncertain significance” (VUS), “likely pathogenic”, and “pathogenic”. Interpreting loss-of-function variants is relatively straightforward, but assessing missense variants poses a challenge. Applying the ACMG criteria frequently leads to the classification of VUS, offering limited value to patients and their clinical management. In ClinVar[20], 92% of missense variants (1902,58 out of 2057,26) were categorized as VUS or variants with inconsistent pathogenicity interpretations[21]. The complexity of categorizing missense variants is further compounded by the prevalence of rare missense variants within individual genomes[22]. While multiplexed assays of variant effect (MAVEs) systematically measure the effects of protein variants[23] and can accurately predict the clinical outcomes of these variants[24], a comprehensive proteome-wide assessment of variant pathogenicity is still lacking due to the cost and labor involved in conducting MAVE experiments. Hence, there arises a necessity for further methods to enhance the classification of missense variants.

Computational methods can utilize patterns in biological data to predict the pathogenicity of unannotated variants, thereby bridging the interpretation gap of these variants. In recent years, a variety of prediction methods have been proposed, and these methods can be divided into two stages with the emergence of the AlphaFold[25]. The methods in the first stage do not rely on the predicted 3D structure by AlphaFold. Representative machine learning-based methods include CADD[26, 27], REVEL[28], and DEOGEN2[29]. Deep learning models can learn high-order dependencies between amino acids from protein sequences and have shown good performance, such as the supervised deep learning methods MVP[30], gMVP[31], and the unsupervised deep learning method EVE[32]. The significant success in predicting protein 3D structures has ushered in a new era for predicting the pathogenicity of missense mutations. AlphaMissense[33], ESM1b[34] has transferred protein 3D structure prediction models to pathogenicity prediction. Recently, with the continuous improvement of AlphaFold database[25, 35], methods based on the predicted structures of AlphaFold have been gradually developed. These methods mainly include SPRI[36], AlphaScore[21], primateAI-3D[37], and others.

In this study, we systematically conducted correlation analysis between allosteric protein mutations and diseases, and performed predictive modeling. We first constructed a dataset of allosteric protein mutations and mapped them to diseases. We found that existing allosteric protein mutations are associated with many diseases, but the clinical significance of the majority of mutations in allosteric proteins remains unclear. Next, by integrating existing machine/ deep learning methods such as CADD, REVEL, DEOGEN2, AlphaScore, gMVP, AlphaMissence and ESM1b, and sequence and structural information before and after amino acid mutations, we developed a meta-method to predict the pathogenicity of mutations at allosteric protein sites. When tested on an independent allosteric protein data set, the proposed method achieved a prediction AUC and AUPR of 0.868 and 0.894, respectively. Specifically, we explored the performance of the proposed and existing methods in predicting the pathogenicity of mutations at allosteric sites. Overall, this study provides insights into exploring the pathogenicity of mutations at allosteric protein offers a powerful tool the identification of pathogenic mutations as well as previously unknown disease-causing allosteric-protein-encoded genes.

## 2 Materials and Methods

### 2.1 Dataset construction

We mined genetic and clinical mutation information from 20,516 human protein-coding genes that have been curated in UniProt[38]. Among these, 20,440 genes contain missense mutation information, and 10,538 genes have mutations related to clinical diseases as recorded in the ClinVar database[25]. 9,790 out of 10,538 genes have predictive results from CADD, REVEL, DEOGEN2, gMVP, ESM1b, AlphaScore, and AlphaMissence, designated as TrainSet.

The list of known human allosteric proteins was downloaded from the Allosteric Database[39] [40] (http://mdl.shsmu.edu.cn/ASD/), totaling 740 protein chains. These proteins are distributed across 90 human genes, referred to as ASDSet. 62 genes in the ASDSet have labels (positive labels are clinical significance recorded as “pathogenic” or “likely pathogenic”; negative labels are clinical significance annotated as “benign” or “likely benign”) in the ClinVar database[25], and with prediction results from CADD, REVEL, DEOGEN2, gMVP, ESM1b, AlphaScore, and AlphaMissence, indicated as TestSet_Gene. The mutations in TestSet_Gene are then mapped to the allosteric-protein-encoded regions (Table S1) and allosteric sites (Table S2) to obtain TestSet_AlloProt and TestSet_AlloSite. TestSet_Gene covers all the labeled mutations on the whole gene sequence, while TestSet_GeneAllo contains the labeled mutations on gene sequence regions aligned with known allosteric protein sequences. TestSet_AlloSite is constructed with labeled mutations on known allosteric sites.

### 2.2 Consensus model based on XGBoost

In this study, three different types of information are used as model input features, including one-hot encoded amino acid mutations, structural-related B-factors and solvent accessibility, as well as predictive values from existing machine learning and deep learning methods such as CADD, REVEL, DEOGEN2, gMVP, ESM1b, AlphaScore, and AlphaMissence. We evaluated more than 50 methods in the field and selected the existing methods as features following the criteria below: the selected methods should have as many predictions as possible within the whole genome; with lower Jaccard similarity coefficient of prediction results between each other; and should be representative or state-of-the-art methods. A Detailed description of these features is listed in Table 1.

**Table 1.** Descriptions of features in MetaMutPre.

| Feature | Dimension | Description |
| --- | --- | --- |
| One-hot encoding | 40 | One-hot encoding of amino acids before and after a site mutation |
| B-factor | 1 | The value that can be used to identify the mobility and flexibility of atoms, amino acid side chains, and loop regions in protein structures[43] |
| Solvent accessibility | 1 | The key feature of proteins for determining their folding and stability[44] |
| CADD[27] | 1 | A logistic regression model trained with over 60 annotations. |
| REVEL[28] | 1 | A random forest based ensemble method incorporating 18 individual pathogenicity prediction scores from 13 tools as predictive features |
| DEOGEN2[29] | 1 | A random forest based model trained with 11 features such as early folding predictions and protein domain-oriented features. |
| gMVP | 1 | A graph attention neural network model trained with features extracted from multi-sequence alignment, gnomAD and predicted structural properties |
| AlphaMissense | 1 | A two-stage model with an adaptation of AlphaFold fine-tuned on human and primate variant population frequency databases |
| ESM1b | 1 | A fine-tune protein language model based on evolutionary features extracted from multi-sequence alignment |
| AlphaScore | 1 | A random forest based model trained on features extracted from protein structures extracted from AlphaFold2 |

The extreme gradient boosting algorithm (XGBoost) is a tree-based prediction model known for its robustness and interpretability, and it is well-suited for outliers, continuous, and categorical variables. During the model training process, we compared various algorithms such as linear regression, SVM, random forest, AdaBoost, etc. However, XGBoost demonstrated the highest overall predictive performance, and is therefore used to construct the pathogenic mutation prediction model. Due to the high number of parameters in XGBoost and the significant impact of different parameter settings on model performance, we used grid search (Table S1) based on cross-validation of TrainSet to determine the best parameter combinations. The selected parameter values for the optimal model are as follows: ‘colsample_bytree’= 0.75, ‘eta’ = 0.1, ‘gamma’ = 0.60, ‘max_depth’ = 3, ‘min_child_weight’ = 4.0, ‘n_estimators’ = 1000.0, ‘subsample’= 0.75; other parameters were set to default values. The model was trained based on TrainSet that is completely independent of TestSet_Gene, TestSet_AlloProt and TestAlloSite.

### 2.3 Evaluation metrics

In order to assess the proposed model, we employ various metrics to evaluate the model’s performance. The key evaluation metrics include area under the curve-receiver operating characteristics (AUC-ROC), and area under precision-recall curve (AUPR), precision, sensitivity (recall), specificity, accuracy, F1 score and Matthews correlation coefficient (MCC), defined as follows.

AUC measures the overall performance of the model at different thresholds. A higher AUC value closer to 1 indicates better model performance and the ability to differentiate between positive and negative samples. AUCPR evaluates how well the model balances precision and recall (sensitivity) at different classification thresholds.

Precision refers to the proportion of truly pathogenic mutations among all samples predicted by the model to be pathogenic mutations, as shown in Eq. (1).

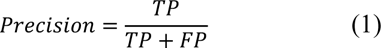

where, *TP* and *FP* represent the number of correctly predicted pathogenic mutations and incorrectly predicted pathogenic mutations in the entire test set, respectively.

Sensitivity (recall), shown in Eq. (2), refers to the proportion of truly pathogenic mutations among all samples that are actually pathogenic mutations and are correctly predicted as such by the model.

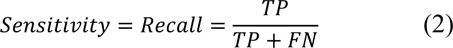

where, *FN* represents the number of benign mutations incorrectly predicted.

Specificity, as in Eq. (3), refers to the proportion of benign mutation samples that are correctly predicted as benign mutations by the model among all samples that are actually benign mutations.

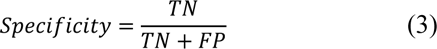

where *TN* is the number of correctly predicted benign mutations.

F1_score, calculated using Eq. (4), represents the harmonic mean of precision and recall (sensitivity), providing a balanced consideration of both precision and recall.

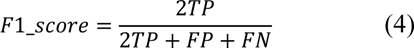

Accuracy, calculated as Eq. (5), refers to the proportion of correctly predicted mutation samples among the total mutation samples.

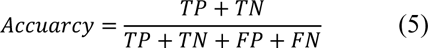

Matthews correlation coefficient (MCC) in Eq. (6) evaluates all four categories in the confusion matrix comprehensively. It is an evaluation metric that can be used regardless of class balance or imbalance, with a range of [-1, 1].

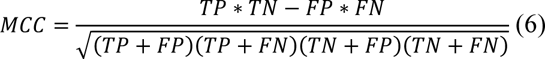

## 3 Results and Discussion

### 3.1 The existing allosteric proteins exhibit high sequence similarities

We first utilized the Needle-Wunsch algorithm in Biopython[41] to perform pairwise sequence alignment for the 740 protein chains in the ASDSet, and then conducted clustering analysis on the similarity of these sequences using the “average linkage” hierarchical clustering algorithm, as shown in **Fig. 1**. It can be observed that many allosteric proteins exhibit high sequence similarity, with protein chains sharing high similarity often originating from the same protein family encoded by the same gene. Conversely, protein chains located on different genes display relatively lower sequence similarity, consistent with the results of sequence identities between genes in **Fig. 2**. **Figure 3** displays the number of allosteric protein chains retained on each gene at different sequence similarity thresholds. For example, at a sequence similarity threshold of 100%, there are 126, 48, 34, 33, 30, and 25 allosteric protein chains on TTR, MAPK14, AMY2A, KIF11, PYGL, and MAP2K1, respectively. When the sequence similarity threshold decreases to 98% and 90%, the numbers of allosteric protein chains on TTR, MAPK14, AMY2A, KIF11, PYGL, and MAP2K1 decrease to 7/ 4/ 1/ 3/ 2/ 8 and 3/ 1/ 1/ 1/ 1/ 4, respectively. The high sequence similarity of allosteric proteins suggests that there are still many undiscovered allosteric proteins distributed across different protein families.

**Fig. 1.**
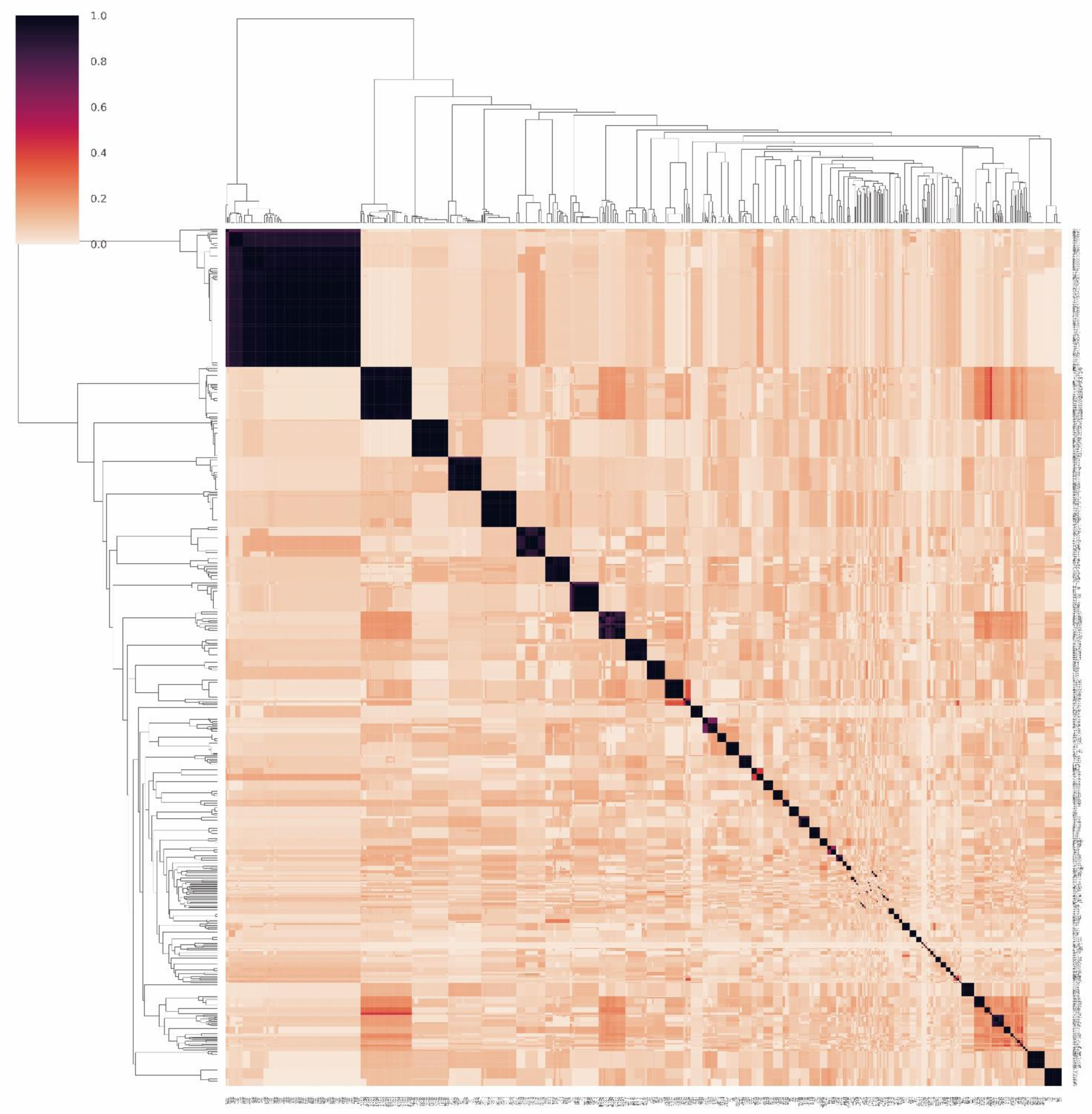
The clustering heatmap of the pairwise protein sequence identities.

**Fig. 2.**
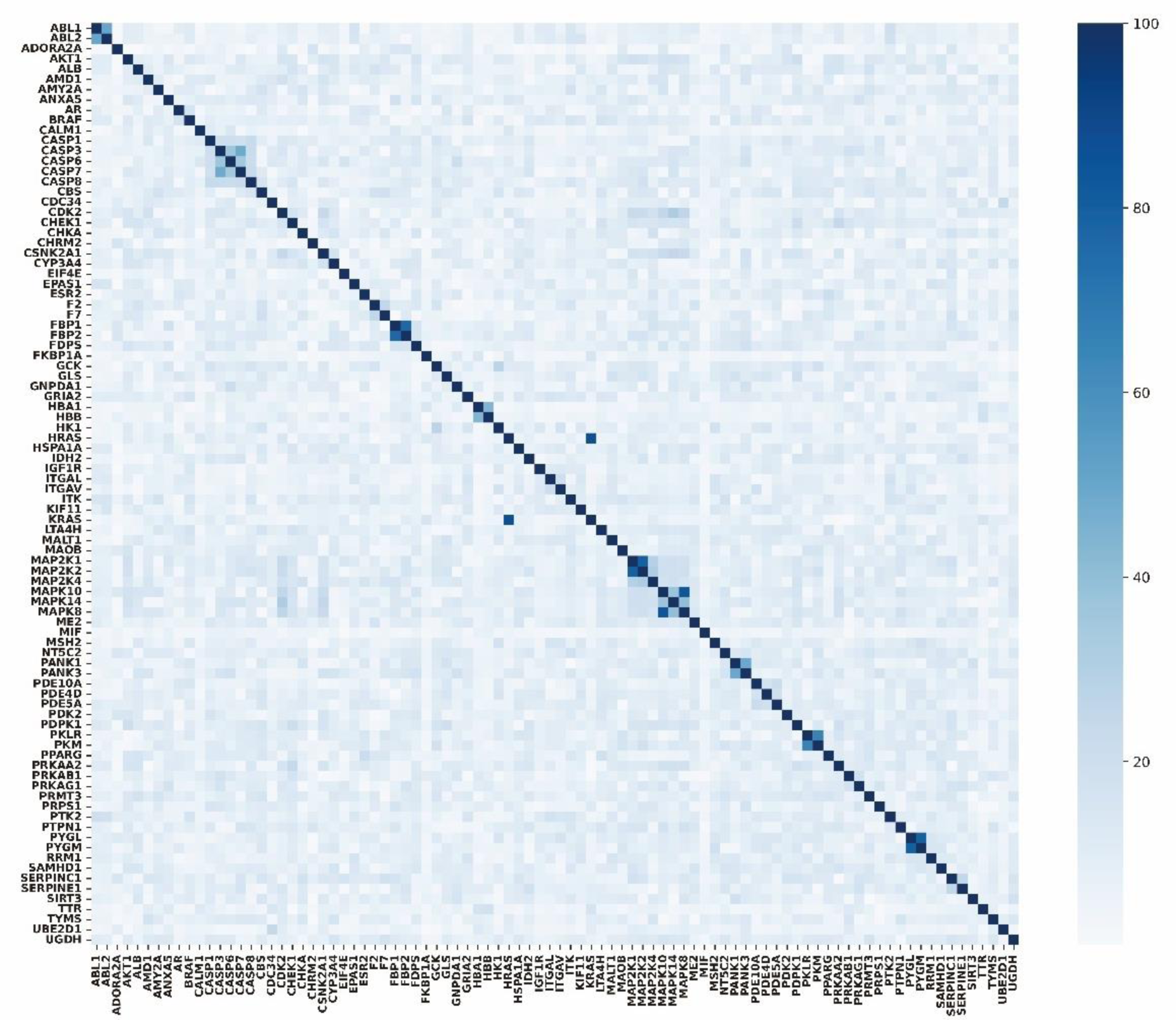
The heatmap of sequence identities between genes.

**Fig. 3.**
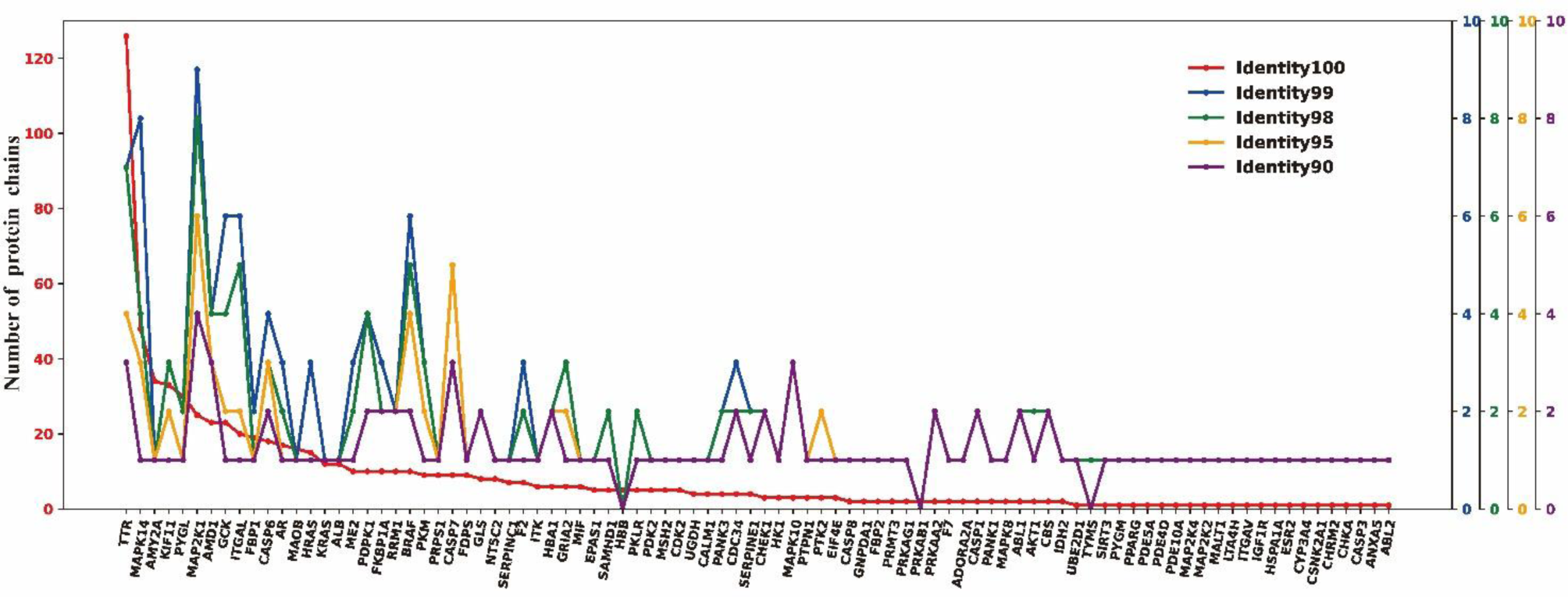
The number of protein chains in each gene with different sequence identity thresholds.

Repeatedly calculating and analyzing allosteric proteins with high sequence similarity would result in significant data redundancy. Therefore, in the subsequent analysis, the corresponding gene name is used to represent the common sequences in each protein cluster.

### 3.2 The clinical significances of more mutations in allosteric proteins await further exploration

As shown in **Fig. S1**, mutations were observed in all genes encoding allosteric proteins, except for PANK1. Among the 89 genes, 47 genes contain clinically known pathogenic mutations, while 64 genes contain benign mutations, and 61 genes harbor mutations with unclear clinical significance. Additionally, 42 genes record both pathogenic and benign mutations, and 41 genes contain mutations of pathogenic, benign, and unclear clinical significance. Subsequently, we analyzed and explored the relationships between the 47 pathogenic mutation-carrying genes and existing diseases (Mutation related disease names were obtained from UniProt https://www.uniprot.org/diseases/ and UniProtKB[42]), as shown in **Fig. 4**, **Fig. 5**, and **Fig. 6**. The phylogenetic trees in the figures represent allosteric proteins encoded by the corresponding gene, and the texts inside the light purple circles indicating the abbreviations of diseases caused by allosteric protein mutations. The disease descriptions can be found in **Table S4**. These allosteric proteins harboring pathogenic mutations that are associated with 115 different diseases.

**Fig. 4.**
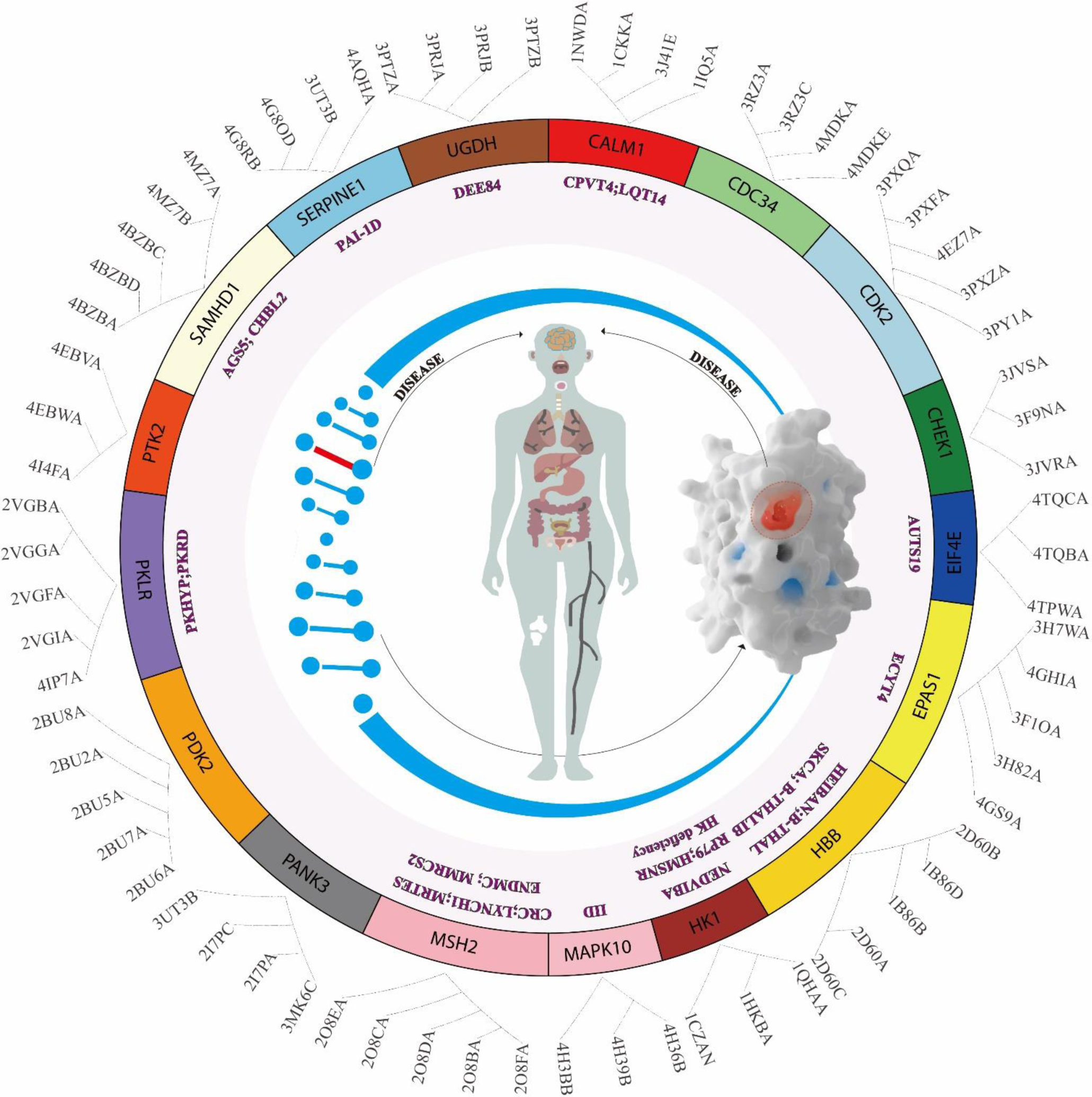
Human diseases caused by pathogenic mutations in known allosteric proteins.

**Fig. 5.**
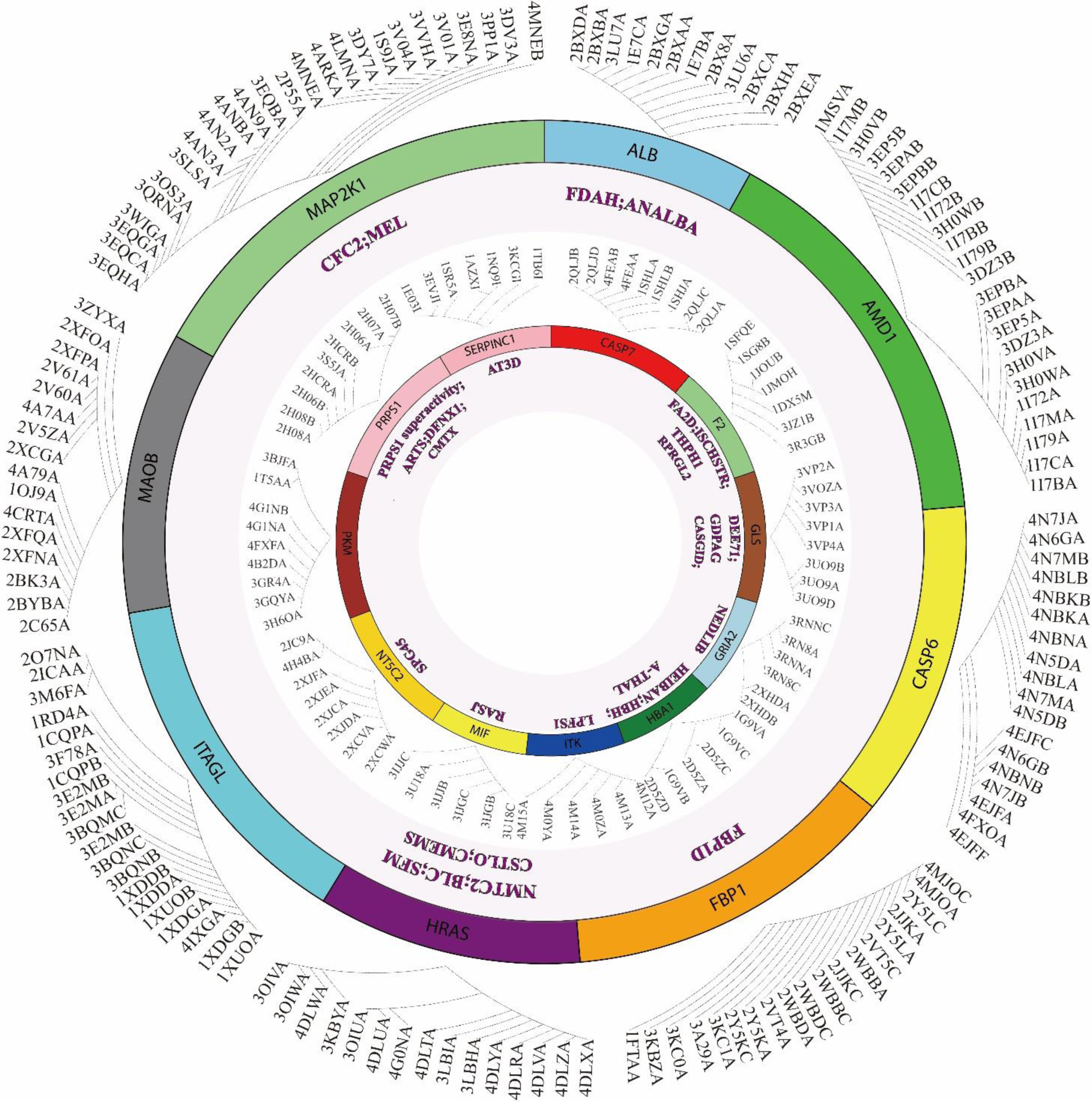
Human diseases caused by pathogenic mutations in known allosteric proteins(continued).

**Fig. 6.**
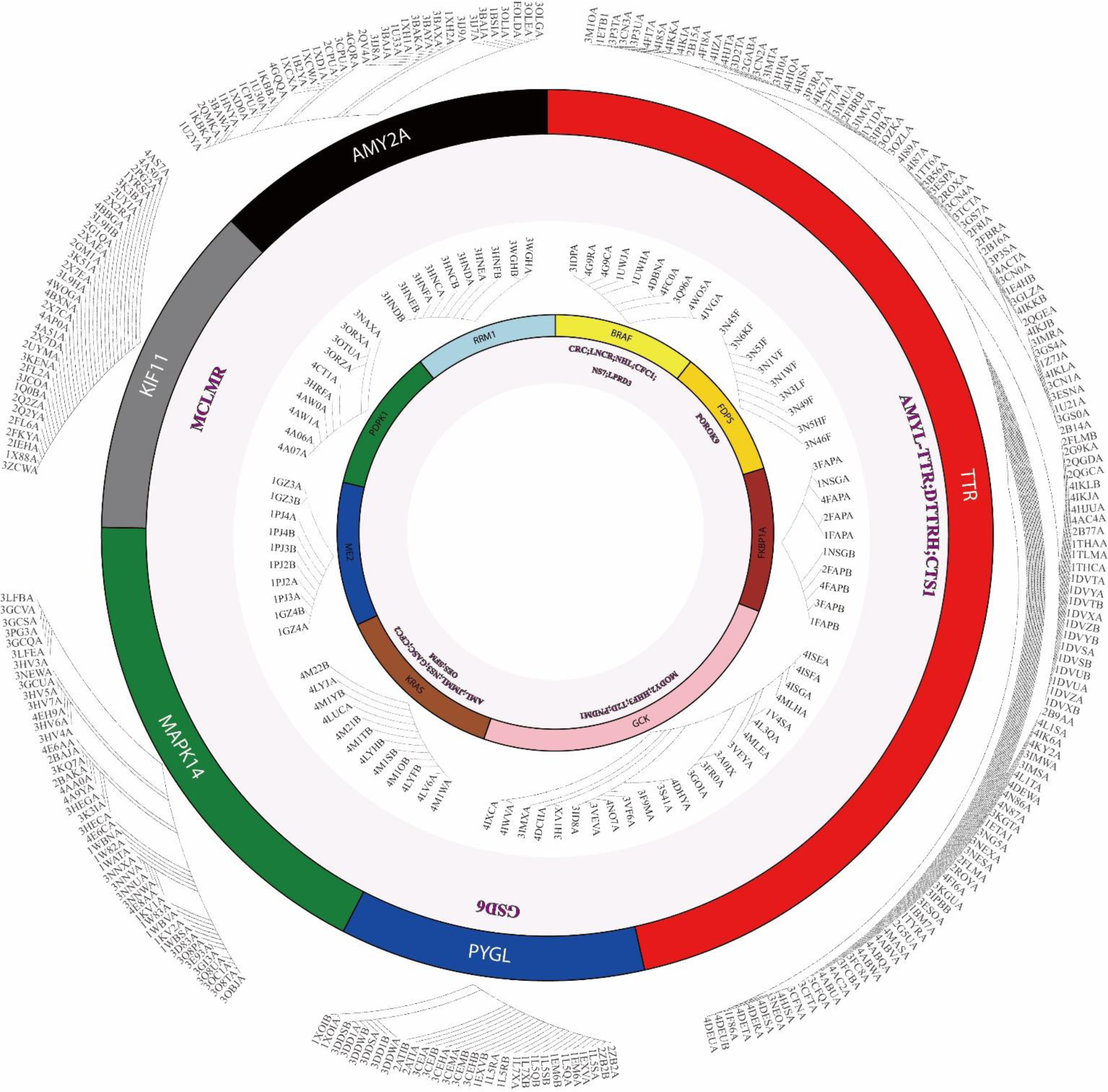
Human diseases caused by pathogenic mutations in known allosteric proteins(continued).

Although mutations in allosteric proteins are closely linked to diseases, the clinical significance of the majority of mutations remains unclear. **Figure 7** documents the number of clinically observed mutations, pathogenic mutations recorded in ClinVar, benign mutations, and mutations of unclear clinical significance in genes encoding allosteric proteins. From **Fig. 6**, it can be seen that the number of pathogenic and benign mutations recorded in databases such as ClinVar is much lower than the number of mutations clinically observed. While mutations in allosteric proteins are intricately linked to diseases, the clinical significance of more mutations awaits further exploration.

**Fig. 7.**
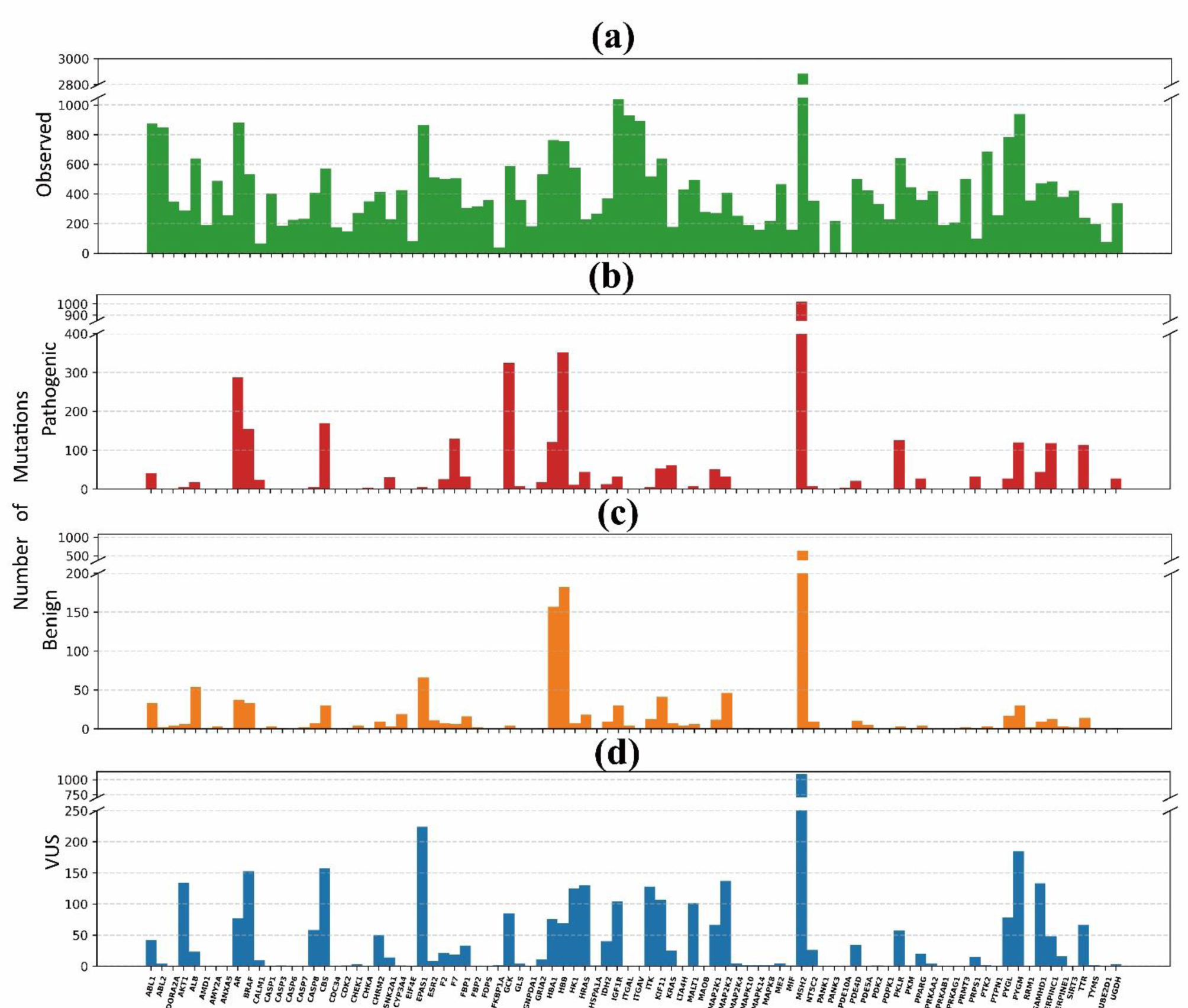
The number of (a) observed mutations; (b) pathogenic mutations; (c) benign mutations; and (d) variants of uncertain significance.

### 3.3 Prediction of pathogenic mutations for allosteric proteins

In order to more effectively explore and analyze the clinical significance of mutations in allosteric proteins, we first analyzed the performance of MetaMutPre in the entire TestSet_Gene dataset and compared it with other methods in terms of AUC, AUPR, MCC, Accuracy, F1 score, Precision, Sensitivity, and Specificity.

Figure 8 and Fig. 9 respectively demonstrate the AUCs of MetaMutPre on the TestSet_Gene and TestSet_GeneAllo. MetaMutPre achieved an AUC of 0.879 on the TestSet_Gene, which is 0.033 higher than the second-ranking gMVP and outperformed REVEL, AlphaMissence, CADD, ESM1b, SPRI, DEOGEN2, and AlphaScore by 0.063, 0.079, 0.115, 0.117, 0.139, 0.175, and 0.186, respectively. MetaMutPre also achieved the highest AUC and AUPR among all prediction methods, The AUC and AUPR of MetaMutPre are 0.036/ 0.073/ 0.085/ 0.124/ 0.148/ 0.13/ 0.164/ 0.182 and 0.031/ 0.061/ 0.063/ 0.132/ 0.128/ 0.12/ 0.086/ 0.143 higher than gMVP/ REVEL/ AlphaMissence/ CADD/ ESM1b/ SPRI/ DEOGEN2/ AlphaScore, respectively.

**Fig. 8.**
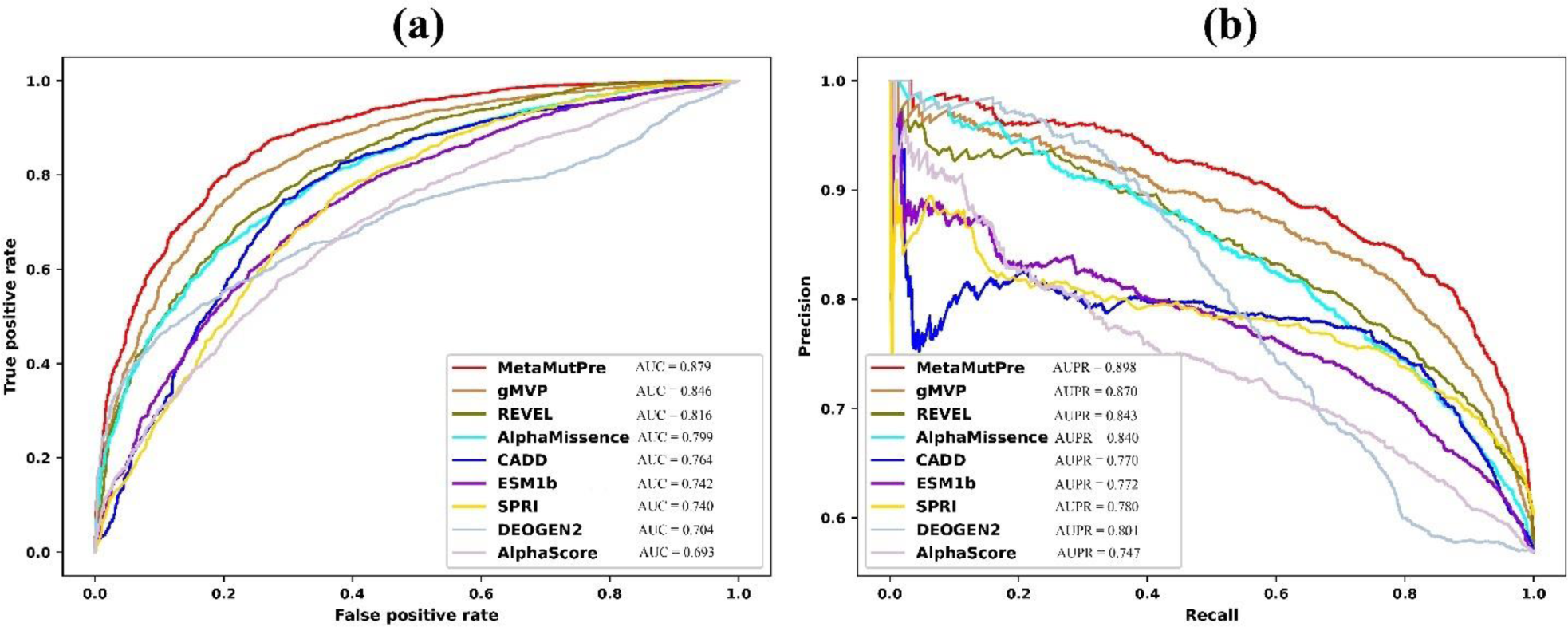
The prediction AUC of MetaMutPre and other methods on TestSet_Gene.

**Fig. 9.**
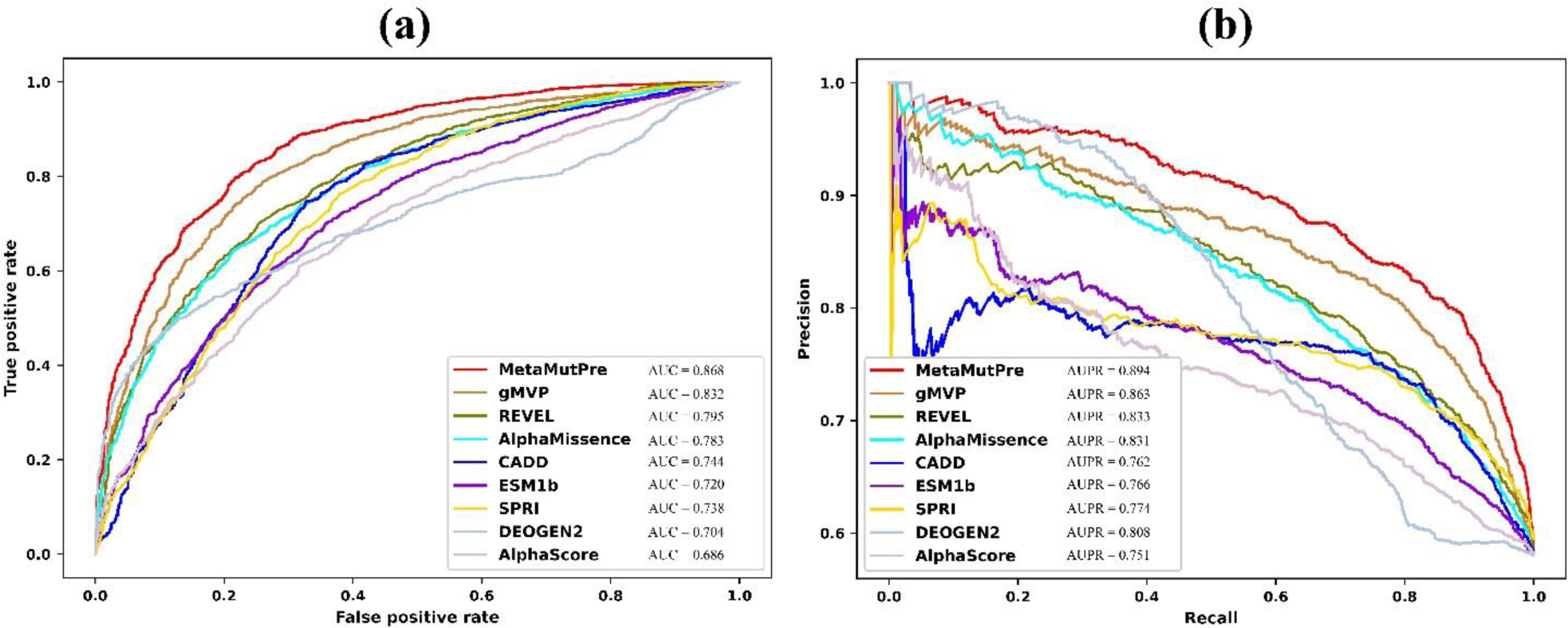
The prediction AUC of MetaMutPre and other methods on TestSet_GeneAllo.

Different models provide different predictive scores for the same mutation. **Table S5** lists the maximum, minimum, median, and mean prediction scores of different methods on the test set and training set. For the predictive performance of the same model, the AUC value is often fixed, but the selection of the threshold has some impact on the evaluation of MCC, Accuracy, F1 score, Precision, Sensitivity, and Specificity. **Table S6** provides the optimal threshold corresponding to the youden_index of the model on the test set. **Table 2** and **Table 3** show the predictive performance of the MetaMutPre and eight other methods based on the youden-index corresponding threshold on TestSet_Gene and TestSet_AlloProt. In terms of MCC, Accuracy, F1 score, and Sensitivity, MetaMutPre is respectively higher than gMVP/ REVEL/ AlphaMissence/ CADD/ ESM1b/ SPRI/ DEOGEN2/ AlphaScore by 0.59/ 0.134/ 0.158/ 0.155/ 0.233/ 0.224/ 0.225/ 0.315, 0.032 /0.067/0.094 /0.078 /0.117 /0.103 / 0.156/ 0.157, 0.035/ 0.063/ 0.114/ 0.075/ 0.11/ 0.078/ 0.223/ 0.145, and 0.066/ 0.083/ 0.203/ 0.1/ 0.136/ 0.09/ 0.369/ 0.168, respectively. In terms of Precision, MetaMutPre surpasses all methods except DEOGEN2, while in Specificity, it is slightly lower than gMVP, AlphaMissence, and DEOGEN2. From Table 3, it can be seen that MetaMutPre outperforms other methods in terms of AUC, MCC, Accuracy, F1 score, and Sensitivity, but lags behind DEOGEN2 in Specificity.

**Table 2.** Performance evaluation of MetaMutPre against other methods on TestSet_Gene. (The threshold is determined by youden index)

| Method | MCC | Accuracy | F1_score | Precision | Sensitivity | Specificity |
| --- | --- | --- | --- | --- | --- | --- |
| MetaMutPre | **0.607** | **0.808** | **0.833** | 0.821 | **0.846** | 0.757 |
| gMVP | 0.548 | 0.776 | 0.798 | 0.818 | 0.78 | 0.771 |
| REVEL | 0.473 | 0.741 | 0.77 | 0.777 | 0.763 | 0.711 |
| AlphaMissence | 0.449 | 0.714 | 0.719 | 0.815 | 0.643 | 0.808 |
| CADD | 0.452 | 0.73 | 0.758 | 0.771 | 0.746 | 0.707 |
| ESM1b | 0.374 | 0.691 | 0.723 | 0.737 | 0.71 | 0.665 |
| SPRI | 0.383 | 0.705 | 0.755 | 0.754 | 0.756 | 0.627 |
| DEOGEN2 | 0.382 | 0.652 | 0.61 | **0.844** | 0.477 | **0.883** |
| AlphaScore | 0.292 | 0.651 | 0.688 | 0.699 | 0.678 | 0.615 |

**Table 3.** Performance evaluation of MetaMutPre against other methods on TestSet_AlloProt. (The threshold is determined by youden index)

| Method | MCC | Accuracy | F1_score | Precision | Sensitivity | Specificity |
| --- | --- | --- | --- | --- | --- | --- |
| MetaMutPre | **0.575** | **0.791** | **0.815** | **0.835** | **0.796** | 0.783 |
| gMVP | 0.527 | 0.768 | 0.798 | 0.808 | 0.789 | 0.74 |
| REVEL | 0.44 | 0.719 | 0.743 | 0.792 | 0.7 | 0.746 |
| AlphaMissence | 0.424 | 0.703 | 0.716 | 0.805 | 0.645 | 0.783 |
| CADD | 0.418 | 0.715 | 0.753 | 0.76 | 0.746 | 0.673 |
| ESM1b | 0.341 | 0.676 | 0.715 | 0.731 | 0.7 | 0.643 |
| SPRI | 0.383 | 0.703 | 0.753 | 0.746 | 0.76 | 0.621 |
| DEOGEN2 | 0.388 | 0.652 | 0.616 | **0.835** | 0.481 | **0.868** |
| AlphaScore | 0.289 | 0.641 | 0.663 | 0.727 | 0.61 | 0.683 |

Comparing the results in **Table 2** and **Table 3**, the performance of MetaMutPre on the TestSet_AlloProt and TestSet_Gene shows, with a difference of only 0.011 in AUC, indicating its suitability for predicting the pathogenicity of mutations in allosteric proteins. Additionally, we can observe that the overall predictive performance of all methods on the TestSet_AlloProt dataset is slightly lower than that on the TestSet_Gene dataset. There may be two possible reasons for this: first, there may not be enough allosteric proteins included in the training set (although the training set may cover many unknown homologous proteins, this cannot be verified at present); second, allosteric proteins, especially the unique physical and chemical properties of homologous regulatory regions, make mutations occurring in these regions more challenging to predict than mutations occurring in regular protein regions.

### 3.4 Exploration of pathogenic mutations on allosteric sites

The allosteric sites remotely control the regulation of many protein activities. The dysfunction of allosteric sites may have significant effects on mediating human diseases.

In this section, we assessed the predictive performance of MetaMutPre and other methods on allosteric sites. As shown in Fig. 10, MetaMutPre exhibits superior AUC and AUPR values compared to other methods, surpassing gMVP/ REVEL/ AlphaMissence/ CADD/ ESM1b/ SPRI/ DEOGEN2/ AlphaScore by 0.029/ 0.06/ 0.037/ 0.078/ 0.148/ 0.218/ 0.034/ 0.118 and 0.011/ 0.035/ 0.012/ 0.026/ 0.053/ 0.098/ 0.005/ 0.035, respectively. demonstrating its ability to strike a balance in recognizing pathogenic and benign mutations. The prediction results in Fig. 11 also demonstrate that there is a trade-off between sensitivity and specificity, and methods based on structural features, such as SPRI and AlphaScore, usually show stronger ability in recognizing benign mutations. Comparing the results on TestSet_Gene and TestSet_AlloProt, it experiences some degradation in overall predictive performance. This may primarily be attributed to two factors. Firstly, the significantly higher number of pathogenic mutations at known allosteric sites compared to benign mutations (226: 39), making benign mutation detection more challenging. Secondly, the distinct physical and chemical properties of allosteric sites compared to non-allosteric sites on proteins, leading to difficulties in predicting mutations occurring on these sites. Nevertheless, MetaMutPre continues to show overall balanced predictions than other methods. Using the threshold (0.941) determined by Youden index, we identified 1764 potential pathogenic mutations on allosteric sites from existing allosteric sites, as shown in Table S7. Among these, 128 out of the 1764 potential pathogenic mutations on allosteric sites have been confirmed by ClinVar.

**Fig. 10.**
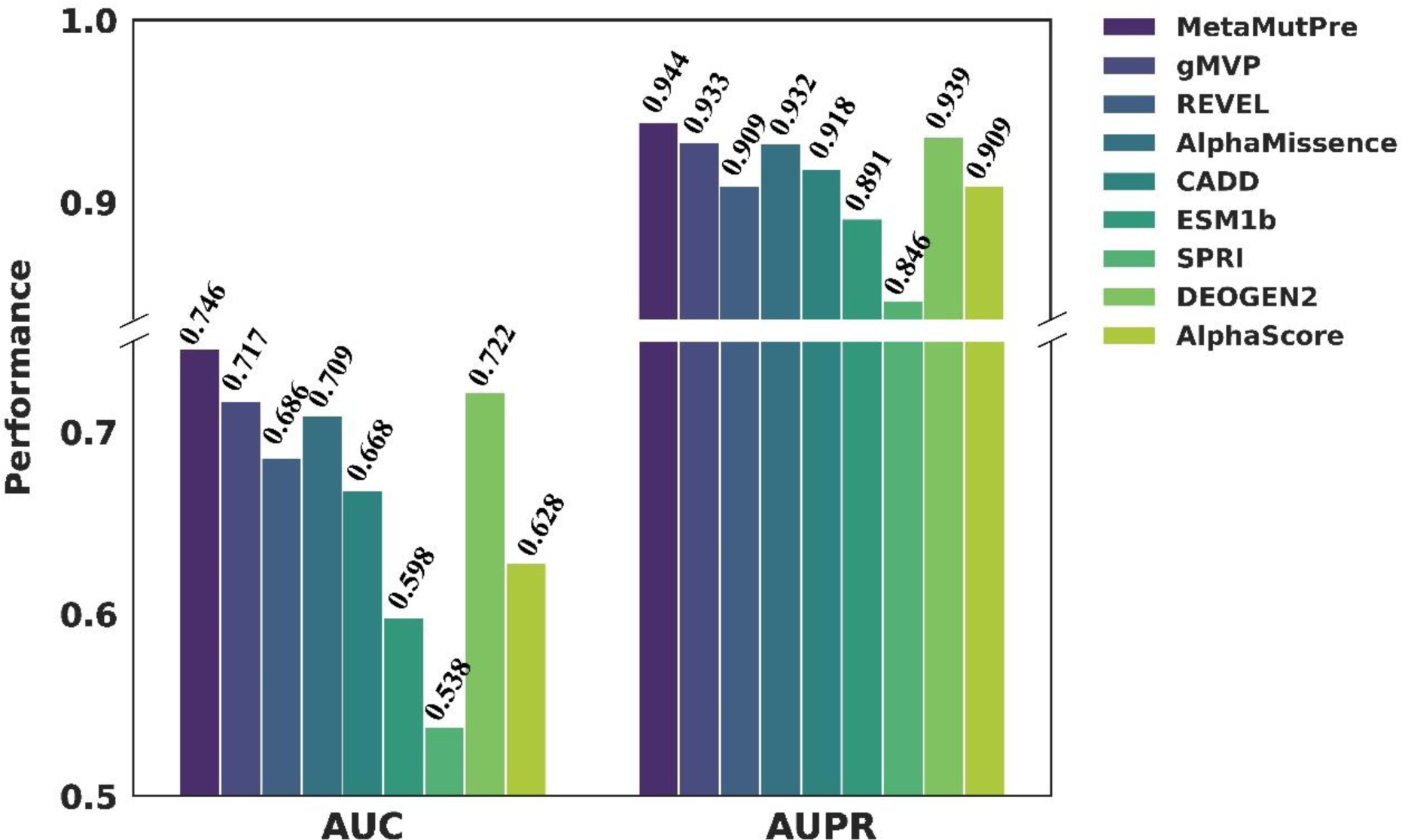
The prediction AUC and AUPR of different methods on allosteric sites.

**Fig 11.**
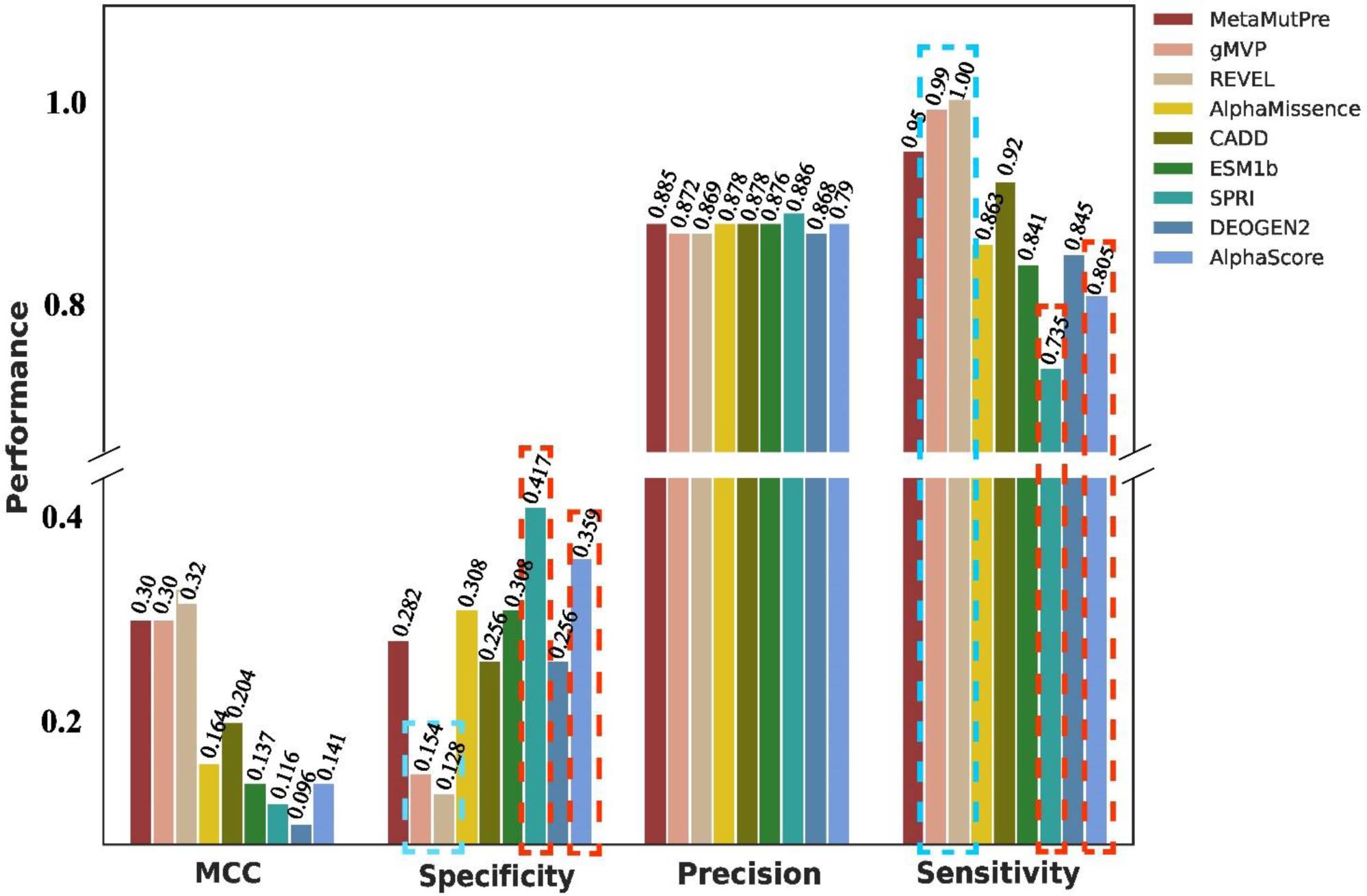
The prediction MCCs, specificities, precisions and sensitivities of different methods on allosteric sites.

## 4 Conclusion

Allosteric regulation is an inherent function of proteins in various physiological and pathological conditions. In this study, we analyzed the mutations in known human allosteric protein, and found that existing allosteric protein mutations are associated with many diseases, but the clinical significance of the majority of mutations remains unclear. Next, we developed an ensemble learning model based on XGBoost for pathogenicity mutation prediction of allosteric proteins. When tested on the benchmark dataset, the proposed method has a higher overall predictive AUC performance on allosteric proteins than the AUC prediction values on allosteric sites. The performance loss on TestSet_AlloSite mainly comes from the inaccuracy in recognizing negative (benign) samples. Methods based on structural features excel in identifying negative samples but face a trade-off between sensitivity and specificity. In predicting the pathogenicity of mutations on allosteric proteins, especially at allosteric sites, we typically need to consider the following challenges. Firstly, the significantly higher number of pathogenic mutations at known allosteric sites compared to benign mutations, making benign mutation detection more challenging. Secondly, the distinct physical and chemical properties of allosteric sites compared to non-allosteric sites on proteins, leading to difficulties in predicting mutations occurring on these sites. Overall, for the pathogenicity prediction for mutations on both allosteric proteins and allosteric sites, MetaMutPre demonstrates a more outstanding and balanced prediction performance towards various metrics.

In summary, the findings in this work illuminate the significance of allosteric mutation in disease processes, and MetaMutPre can serve as a valuable tool for the identification of pathogenic allosteric mutations as well as previously unknown disease-causing genes.

