## Supplementary material for "Exploring Pathogenic Mutation in Allosteric Proteins: the Prediction and Beyond": Suppl

### [Supplementary Files]

Huiling Zhang[1], Zhen Ju[2], Jingjing Zhang[2], Xijian Li[1], Hanyang Xiao[1], Xiaochuan Chen[1], Yuetong li[1], Xinran Wang[1], and Yanjie Wei[2,*]

[1] College of Mathematics and Information, South China Agriculture University, Guangzhou 510640, China.
[2] Shenzhen Institute of Advanced Technology, Chinese Academy of Sciences, Shenzhen 518055, China.

**This supplementary file contains**

### Figures

**Fig. S1 The allosteric protein encoded genes with known pathogenic, benign, uncertain and observed mutations.**

### Tables

**Table S1 The starting and ending positions of the gene sequences encoding known allosteric proteins**

**Table S2 Known allosteric sites on different genes**

**Table S3 Parameter settings of grid search**

**Table S4 Descriptions of mutation related diseases in allosteric proteins**

**Table S5 The maximum, minimum, median and average output scores of different prediction methods**

**Table S6 Optimal threshold determined by Youden index**

**Table S7 Potential pathogenic mutation on allosteric sites predicted by MetaMutPre**

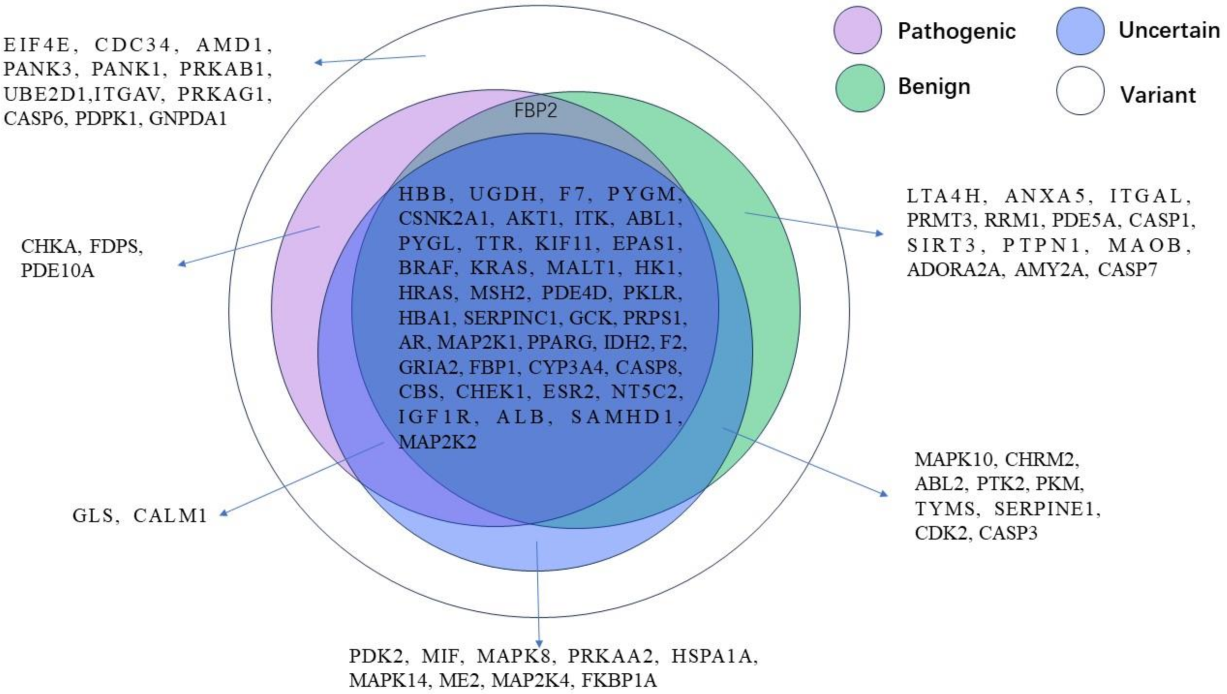


**Fig. S1 The allosteric protein encoded genes with known pathogenic, benign, uncertain and observed mutations**

\

**Table S1. The starting and ending positions of the gene sequences encoding known allosteric proteins**

| Gene | Start | End | Allosteric protein chains |
|---|---|---|---|
| TTR | 1 | 147 | 1BM7A;1DVSA;1DVSB;1DVTA;1DVTB;1DVUA;1DVUB;1DVXA;1DVXB;1DVYA;1DVYB;1DVZA;1DVZB;1E4HB;1ETA1;1ETB1;1F86A;1THAA;1THCA;1TLMA;1TT6A;1TYRA;1U21A;1Y1DA;1Z7JA;2B14A;2B15A;2B16A;2B77A;2B9AA;2F7IA;2F8IA;2FBRA;2FBRB;2FLMA;2FLMB;2G5UA;2G9KA;2GABA;2QGCA;2QGDA;2QGEA;2ROXA;2ROYA;3B56A;3CFNA;3CFQA;3CFTA;3CN0A;3CN1A;3CN2A;3CN3A;3CN4A;3D2TA;3ESNA;3ESOA;3ESPA;3FC8A;3FCBA;3GLZA;3GS0A;3GS4A;3GS7A;3HJ0A;3IMRA;3IMSA;3IMTA;3IMUA;3IMVA;3IMWA;3IPBA;3IPBB;3KGTA;3KGUA;3M1OA;3NEOA;3NESA;3NEXA;3NG5A;3OZKA;3OZLA;3P3RA;3P3SA;3P3TA;3P3UA;3TCTA;4ABQA;4ABUA;4ABVA;4ABWA;4AC2A;4AC4A;4ACTA;4DERA;4DESA;4DETA;4DEUA;4DEUB;4DEWA;4FI6A;4FI7A;4FI8A;4HIQA;4HISA;4HJSA;4HJTA;4HJUA;4I85A;4I87A;4I89A;4IIZA;4IK6A;4IK7A;4IKIA;4IKJA;4IKJB;4IKKA;4IKKB;4IKLA;4IKLB;4KY2A;4L1SA;4L1TA;4MASA;4N86A;4N87A |
| MAPK14 | 1 | 360 | 3K3IA;1WBSA;3KQ7A;3HV6A;3D83A;1WBTA;3E93A;3GCQA;3HV3A;3O8TA;3PG3A;3O8UA;4E6AA;3NNVA;3O8PA;3GCUA;4EH9A;3GCSA;2BAKA;4A9YA;3HEGA;1W82A;3HV5A;2BAJA;3NNXA;3LFEA;3GCVA;4E6CA;3GI3A;3OBJA;1WBNA;3NNUA;3HV7A;1KV1A;1W83A;3HECA;3NEWA;3OC1A;3LFBA;3HV4A;4E8AA;1KV2A;4AA0A;3NNWA;3HV6A;1WBVA |
| AMY2A | 16 | 511 | 1B2YA;1BSIA;1CPUA;1HNYA;1KBBA;1KBKA;1U2YA;1U30A;1U33A;1XCWA;1XCXA;1XD0A;1XD1A;1XH1A;1XH2A;2CPUA;2QMKA;2QV4A;3BAIA;3BAJA;3BAKA;3BAWA;3BAXA;3BAYA;3CPUA;3IJ7A;3IJ8A;3IJ9A;3OLDA;3OLEA;3OLGA;3OLIA;4GQQA;4GQRA |
| KIF11 | 1 | 369 | 1Q0BA;1X88A;1YRSA;2FKYA;2FL2A;2FL6A;2G1QA;2GM1A;2IEHA;2PG2A;2Q2YA;2Q2ZA;2UYIA;2UYMA;2WOGA;2X2RA;2X7CA;2X7DA;2X7EA;2XAEA;3CJOA;3K3BA;3K5EA;3KENA;3L9HA;3L9HB;3ZCWA;4A50A;4A51A;4AP0A;4AS7A;4BBGA;4BXNA |
| PYGL | 1 | 847 | 1EM6A;1EM6B;1EXVA;1EXVB;1L5QA;1L5QB;1L5RA;1L5RB;1L5SA;1L5SB;1L7XA;1L7XB;1XOIA;1XOIB;2ATIA;2ATIB;2ZB2A;2ZB2B;3CEHA;3CEHB;3CEJA;3CEJB;3CEMA;3CEM |

| | | | |
|---|---|---|---|
| | | | B;3DD1A;3DD1B;3DDSA;3DDSB;3DDWA;3DDWB |
| MAP2K1 | 34 | 393 | 1S9JA;2P55A;3DV3A;3DY7A;3E8NA;3EQBA;3EQCA;3EQGA;3EQHA;3ORNA;3OS3A;3PP1A;3SLSA;3V01A;3V04A;3VVHA;3WIGA;4AN2A;4AN3A;4AN9A;4ANBA;4ARKA;4LMNA;4MNEA;4MNEB |
| AMD1 | 1 | 334 | 1I72A;1I72B;1I79A;1I79B;1I7BA;1I7BB;1I7CA;1I7CB;1I7MA;1I7MB;1MSVA;3DZ3A;3DZ3B;3EP5A;3EP5B;3EPAA;3EPAB;3EPBA;3EPBB;3H0VA;3H0VB;3H0WA;3H0WB |
| GCK | 1 | 465 | 1V4SA;3A0IX;3F9MA;3FR0A;3GOIA;3H1VX;3ID8A;3IMXA;3S41A;3VEVA;3VEYA;3VF6A;4DCHA;4DHYA;4ISEA;4ISFA;4ISGA;4IWVA;4IXCA;4L3QA;4MLEA;4MLHA;4NO7A |
| ITGAL | 146 | 336 | 1CQPA;1CQPB;1RD4A;1XDDA;1XDDB;1XDGA;1XDGB;1XUOA;1XUOB;2ICAA;2O7NA;3BQMB;3BQMC;3BQNB;3BQNC;3E2MA;3E2MB;3F78A;3M6FA;4IXDA |
| FBP1 | 1 | 338 | 1FTAA;2JJKA;2JJKC;2VT5A;2VT5C;2WBBA;2WBBC;2WBDA;2WBDC;2Y5KA;2Y5KC;2Y5LA;2Y5LC;3A29A;3KBZA;3KC0A;3KC1A;4MJOA;4MJOC |
| CASP6 | 1 | 292 | 4EJFA;4EJFC;4EJFF;4FXOA;4N5DA;4N5DB;4N6GA;4N6GB;4N7JA;4N7JB;4N7MA;4N7MB;4NBKA;4NBKB;4NBLA;4NBLB;4NBNA;4NBNB |
| AR | 665 | 901 | 2PIOA;2PIPL;2PIQA;2PITA;2PIUA;2PIVA;2PIWA;2PIXA;2PKLA;2YHDA;2YLOA;2YLPA;2YLQA;3ZQTA;4HLWA;4K7AA |
| MAOB | 1 | 520 | 1OJ9A;2BK3A;2BYBA;2C65A;2V5ZA;2V60A;2V61A;2XCGA;2XFNA;2XFOA;2XFPA;2XFQA;3ZYXA;4A79A;4A7AA;4CRTA |
| HRAS | 1 | 166 | 3K8YA;3LBHA;3LBIA;3OIUA;3OIVA;3OIWA;4DLRA;4DLTA;4DLUA;4DLVA;4DLWA;4DLXA;4DLYA;4DLZA;4G0NA |
| KRAS | 1 | 169 | 4LUCA;4LV6A;4LYFB;4LYHB;4LYJA;4M1OB;4M1SB;4M1TB;4M1WA;4M1YB;4M21B;4M22B |
| ALB | 25 | 609 | 1E7BA;1E7CA;2BX8A;2BXAA;2BXBA;2BXCA;2BXDA;2BXEA;2BXGA;2BXHA;3LU6A;3LU7A |

| | | | |
|---|---|---|---|
| ME2 | 21 | 584 | 1GZ3A;1GZ3B;1GZ4A;1GZ4B;1PJ2A;1PJ2B;1PJ3A;1PJ3B;1PJ4A;1PJ4B |
| PDPK1 | 44 | 362 | 3HRFA;3NAXA;3ORXA;3ORZA;3OTUA;4A06A;4A07A;4AW0A;4AW1A;4CT1A |
| FKBP1A | 2 | 108 | 1FAPA;1FAPB;1NSGA;1NSGB;2FAPA;2FAPB;3FAPA;3FAPB;4FAPA;4FAPB |
| RRM1 | 1 | 792 | 2WGHA;2WGHB;3HNCA;3HNCB;3HNDA;3HNDB;3HNEA;3HNEB;3HNFA;3HNFB |
| BRAF | 421 | 728 | 1UWHA;1UWJA;3IDPA;3Q96A;4DBNA;4FC0A;4G9CA;4G9RA;4JVGA;4WO5A |
| PKM | 1 | 531 | 1T5AA;3BJFA;3GQYA;3GR4A;3H6OA;4B2DA;4FXFA;4G1NA;4G1NB |
| PRPS1 | 1 | 318 | 2H06A;2H06B;2H07A;2H07B;2H08A;2H08B;2HCRA;2HCRB;3S5JA |
| CASP7 | 50 | 303 | 1SHJA;1SHLA;1SHLB;2QLJA;2QLJB;2QLJC;2QLJD;4FEAA;4FEAB |
| FDPS | 70 | 419 | 3N1VF;3N1WF;3N3LF;3N45F;3N46F;3N49F;3N5HF;3N5JF;3N6KF |
| GLS | 71 | 550 | 3UO9A;3UO9B;3UO9D;3VOZA;3VP1A;3VP2A;3VP3A;3VP4A |
| NT5C2 | 1 | 536 | 2JC9A;2XCVA;2XCWA;2XJCA;2XJDA;2XJEA;2XJFA;4H4BA |
| SERPINC1 | 33 | 464 | 1AZXI;1E03I;1NQ9I;1SR5A;1TB6I;3EVJI;3KCGI |
| F2 | 363 | 622 | 1DX5M;1JMOH;1JOUB;1SFQE;1SG8B;3JZ1B;3R3GB |
| ITK | 354 | 620 | 4M0YA;4M0ZA;4M12A;4M13A;4M14A;4M15A |
| HBA1 | 1 | 142 | 1G9VA;1G9VB;1G9VC;2D5ZA;2D5ZC;2D5ZD |
| GRIA2 | 404 | 812 | 2XHDA;2XHDB;3RN8A;3RN8C;3RNNA;3RNNC |
| MIF | 2 | 115 | 3IJGB;3IJGC;3IJJB;3IJJC;3U18A;3U18C |
| EPAS1 | 241 | 350 | 3F1OA;3H7WA;3H82A;4GHIA;4GS9A |
| SAMHD1 | 110 | 626 | 4BZBA;4BZBC;4BZBD;4MZ7A;4MZ7B |

| | | | |
|---|---|---|---|
| HBB | 2 | 147 | 1B86B;1B86D;2D60A;2D60B;2D60C |
| PKLR | 32 | 574 | 2VGBA;2VGFA;2VGGA;2VGIA;4IP7A |
| PDK2 | 14 | 407 | 2BU2A;2BU5A;2BU6A;2BU7A;2BU8A |
| MSH2 | 1 | 934 | 2O8BA;2O8CA;2O8DA;2O8EA;2O8FA |
| CDK2 | 1 | 298 | 3PXFA;3PXQA;3PXZA;3PY1A;4EZ7A |
| UGDH | 1 | 494 | 3PRJA;3PRJB;3PTZA;3PTZB |
| CALM1 | 1 | 149 | 1CKKA;1IQ5A;1NWDA;3J41E |
| PANK3 | 1 | 368 | 2I7PA;2I7PC;3MK6A;3MK6C |
| CDC34 | 2 | 184 | 3RZ3A;3RZ3C;4MDKA;4MDKE |
| SERPINE1 | 20 | 402 | 3UT3B;4AQHA;4G8OD;4G8RB |
| CHEK1 | 2 | 406 | 3F9NA;3JVRA;3JVSA |
| HK1 | 1 | 917 | 1CZAN;1HKBA;1QHAA |
| MAPK10 | 45 | 400 | 4H36A;4H39A;4H3BA |
| PTPN1 | 1 | 300 | 1T49A |
| PTK2 | 411 | 686 | 4EBVA;4EBWA;4I4FA |
| EIF4E | 27 | 217 | 4TPWA;4TQBA;4TQCA |
| CASP8 | 385 | 479 | 3KJNB;3KJQB |
| GNPDA1 | 1 | 289 | 1NE7A;1NE7B |
| FBP2 | 2 | 339 | 3IFAA;3IFCA |
| PRMT3 | 210 | 531 | 3SMQA;4HSGA |
| PRKAG1 | 1 | 331 | 4CFEE;4CFFE |
| PRKAB1 | 1 | 270 | 4CFFA;4CFFB |
| PRKAA2 | 1 | 552 | 4CFEA;4CFEB |
| F7 | 213 | 466 | 1DVAI;1YGCH |
| ADORA2A | 1 | 361 | 3VG9A;3VGAA |
| CASP1 | 122 | 300 | 2FQQA;2FQQB |
| PANK1 | 234 | 593 | 2I7NA;2I7NB |
| MAPK8 | 1 | 370 | 3O2MA;3O2MB |
| ABL1 | 1 | 512 | 1OPLA;3PYYA |
| AKT1 | 1 | 446 | 3O96A;4EJNA |
| CBS | 1 | 551 | 4PCUB;4UUUB |
| IDH2 | 41 | 452 | 4JA8A;4JA8B |
| UBE2D1 | 1 | 147 | 4QPLA |

| | | | |
|---|---|---|---|
| TYMS | 1 | 313 | 2ONBA |
| SIRT3 | 121 | 399 | 4C7BA |
| PYGM | 1 | 842 | 1Z8DA |
| PPARG | 238 | 505 | 3K8SB |
| PDE5A | 526 | 858 | 1TBFA |
| PDE4D | 379 | 794 | 3IADA |
| PDE10A | 515 | 703 | 2ZMFA |
| MAP2K4 | 79 | 399 | 3ALOA |
| MAP2K2 | 59 | 400 | 1S9IA |
| MALT1 | 338 | 720 | 4I1RA |
| LTA4H | 1 | 611 | 3FUDA |
| ITGAV | 12 | 1038 | 1L5GB |
| IGF1R | 983 | 1286 | 3LW0A |
| HSPA1A | 1 | 382 | 4IO8A |
| ESR2 | 257 | 502 | 2FSZB |
| CYP3A4 | 23 | 503 | 1W0FA |
| CSNK2A1 | 1 | 334 | 3H30A |
| CHRM2 | 1 | 466 | 4MQTA |
| CHKA | 75 | 457 | 3ZM9A |
| CASP3 | 180 | 277 | 3KJFB |
| ANXA5 | 1 | 320 | 1HAKA |
| ABL2 | 278 | 546 | 3GVUA |

**Table S2 Known allosteric sites on different genes**

| Gene | Allosteric site |
|---|---|
| BRAF | 462I; 463I; 467S; 470V; 471V; 472Y; 480A; 481A; 482K; 482V; 483K; 497A; 500E; 501E; 503V; 504L; 504V; 505L; 507T; 512I; 513I; 513L; 514L; 515L; 516F; 526I; 527I; 528T; 528V; 529Q; 529T; 530Q; 530W; 531C; 531W; 532C; 532E; 533E; 533G; 535S; 536S; 566L; 567L; 571I; 572I; 573H; 574H; 580N; 581N; 582F; 583F; 591I; 592G; 592I; 593D; 593G; 594D; 594F; 595F; 596G; 597L; 598A; 599T; 601K; 601S |
| CHKA | 117R; 124L; 146R; 153N; 154C; 155D; 156R; 157D; 164M; 165D; 166C; 167E; 168D; 169D; 170E; 171V; 172L; 175E; 176D; 177L; 178Q; 179D; 180M; 194P; 206E; 207Q; 208F; 209I; 212R; 213R; 214F; 252E; 306D; 308Q; 310G; 329I; 330D; 349E; 351E; 354Y; 362K; 364L; 369D; 370S; 371P; 372R; 373N; 374R; 376L; 377K; 380P; 386G; 387P; 388D; 389F; 391Y; 393T; 397Q; 420W; 423W; 435F; 440Y |
| CALM1 | 16F; 17S; 19F; 20D; 21K; 22D; 23G; 24D; 25G; 26T; 27I; 28T; 30K; 31E; 32L; 52I; 53N; 54E; 55V; 56D; 57A; 58D; 59G; 60N; 61G; 62T; 63I; 64D; 65F; 66P; 67E; 68F; 71M; 77K; 86R; 89F; 90R; 92F; 93D; 94K; 95D; 96G; 97N; 98G; 99Y; 100I; 101S; 102A; 103A; 104E; 105L; 125I; 126R; 127E; 128A; 129D; 130I; 131D; 132G; 133D; 134G; 135Q; 136V; 137N; 138Y; 139E; 140E; 141F; 143Q; 144M |
| PYGL | 37F; 38T; 39L; 40V; 41K; 42D; 44N; 45V; 53F; 57H; 60R; 63L; 64V; 67W; 68I; 71Q; 72Q; 75Y; 81R; 155Y; 184R; 185Y; 186G; 187N; 188P; 189W; 190E; 191K; 192S; 193R; 194P; 196F; 226Y; 227D; 229P; 240T; 242R; 282N; 283D; 284N; 285F; 306D; 309R; 310R; 313A; 380L; 382E; 571H; 573Y; 610A; 612G; 613Y; 770R; 771F; 1038T; 1040V; 1053F; 1057H; 1185Y; 1186G; 1187N; 1188P |
| MAPK14 | 30V; 31G; 32S; 33G; 35Y; 38V; 51A; 52V; 53K; 67R; 70R; 71E; 74L; 75L; 78M; 83V; 84I; 104L; 105V; 106T; 107H; 108L; 109M; 110G; 111A; 112D; 115N; 141I; 146I; 147I; 148H; 149R; 150D; 155N; 166I; 167L; 168D; 168G; 169F; 169Y; 170G; 170Y; 171L; 172C; 191P; 192E; 195L; 196N; 197W; 198M; 232L; 242P; 246L; 249K; 250I; 251S; 252S; 255A; 259I; 291L; 292D; 293S; 294D |
| MAP2K1 | 77G; 78N; 79G; 80G; 97K; 99I; 115L; 116L; 118L; 119L; 127V; 128G; 128V; 129F; 129G; 130F; 141I; 142I; 143M; 144M; 188H; 189H; 189R; 190D; 190R; 191D; 192K; 195N; 207C; 208C; 208D; 209D; 209F; 210F; 210G; 211G; 211V; 212S; 212V; 213S; 215L; 216I; 216L; 217I; 219M; 220M; 221N; 222N; 223S; 225G; 226T; 234R; 662R |
| KIF11 | 104Y; 112T; 116E; 117G; 118E; 119R; 127W; 130D; 132L; 133A; 133D; 136I; 137P; 160L; 171L; 172L; 210V; 211Y; 213I; 214L; 215E; 216K; 217G; 218A; 219A; 220K; 221R; 224A; 228M; 229N; 232S; 239F; 266L; 269S; 270E; 287N; 288I; 289N; 292L; 293L; 295L; 296G; 297R; 299I; 300T; 332I; 352Y; 353A; 355R; 356A |
| PDPK1 | 76K; 88L; 91G; 94S; 109A; 111K; 112I; 113L; 113V; 115K; 118I; 118V; 119I; 124I; 124V; 125P; 126Y; 127V; 128Q; 128T; 130E; 131K; 131R; 134M; 142F; 143V; 148C; 148T; 149F; 150Q; 155L; 156Y; 157F; 157L; 159L; 160S; 161Y; 162A; 196L; 201I; 203H; 212L; 221I; 222T; 223D; 224F; 225G; 227A |
| PKLR | 421T; 443L; 444T; 445T; 446T; 447G; 448R; 449S; 450A; 467R; 474L; 475T; 476T; 477T; 478G; 479H; 479R; 480S; 494W; 497D; 498R; 498V; 501R; 524V; 525T; 525W; 526G; 527W; 528R; 529P; 530G; 531S; 532G; 532R; 533Y; 534T; 557G; 559R; 560P; 561G; 562S; 563G; 564Y; 565T |
| ALB | 150Y; 153E; 192S; 196Q; 198L; 199K; 202S; 206F; 209R; 210A; 211F; 212K; 213A; 214W; 216V; 218R; 219L; 222R; 223F; 232S; 235V; 238L; 242H; 257R; 260L; 261A; 264I; 287S; |

| | |
|---|---|
| | 290I; 291A; 292E; 324D; 327L; 328G; 331L; 347L; 350A; 351K; 354E; 480S; 481L; 482V |
| PKM | 26F; 27L; 30M; 43R; 44N; 45T; 46G; 70N; 106R; 311K; 353L; 354D; 389I; 390Y; 393Q; 394L; 397E; 431L; 432T; 433K; 434S; 435G; 436R; 437S; 464H; 468G; 469I; 470F; 471P; 482W; 489R; 514G; 515W; 516R; 517P; 518G; 519S; 520G; 521F; 522T |
| GCK | 61Y; 62V; 63R; 64S; 65T; 66P; 67E; 68G; 69S; 70E; 91V; 96E; 97G; 98Q; 99W; 159I; 207V; 210M; 211I; 212S; 214Y; 215Y; 216E; 218H; 220C; 221E; 235M; 250R; 451L; 452V; 453S; 454A; 455V; 456A; 458K; 459K |
| SAMHD1 | 113D; 114T; 116K; 117V; 118I; 119N; 120D; 125H; 133V; 136I; 137D; 142Q; 145R; 155Y; 156V; 157F; 165F; 169L; 324G; 325I; 330D; 333R; 337F; 352R; 354K; 358N; 372R; 376H; 377K; 378V; 451R; 455K; 523K |
| SERPINE1 | 12A; 37Y; 41S; 44A; 45M; 66I; 75L; 76R; 78L; 79Y; 80K; 93T; 94T; 95D; 116L; 117F; 118R; 119S; 120T; 122K; 139W; 143H; 175W; 176K; 177T; 179F; 204Q; 206N; 207K; 208F; 209N; 226L; 227P; 231D; 268R |
| PDK2 | 23L; 27Q; 28F; 31F; 40T; 41S; 44F; 45L; 53L; 63L; 64P; 66R; 67V; 73V; 80Y; 111I; 112R; 115H; 122M; 125G; 126V; 129Y; 142N; 143I; 146F; 147L; 154R; 157I; 158R; 160L; 161I; 163Q; 164H; 167I; 374Y |
| CASP6 | 36K; 39R; 40L; 41H; 42C; 43V; 44E; 45W; 46T; 47I; 48L; 56F; 57W; 58H; 93A; 94E; 97L; 101H; 125N; 131D; 132A; 133K; 135E; 137Q; 142L; 195A; 196S; 197V; 198Y; 199T; 200L; 201P; 214E; 244E; 287H |
| ITGAL | 128G; 130V; 132L; 134F; 140M; 153F; 157V; 161L; 166Y; 203L; 231T; 233V; 235I; 255I; 257Y; 258I; 259I; 284E; 285F; 286V; 287K; 298L; 301E; 302L; 303Q; 304K; 305K; 306I; 307Y; 308V; 309I |
| RRM1 | 3V; 5K; 6R; 11E; 12R; 13V; 14M; 17K; 18I; 21R; 53T; 56L; 57D; 88K; 226D; 227S; 228I; 229E; 231I; 243K; 255I; 256R; 261Y; 262I; 263A; 264G; 265T; 269S; 270N; 285Y; 286V; 287D; 288Q; 289G |
| AR | 672P; 673I; 674F; 713L; 717V; 718K; 721K; 724P; 725G; 726F; 727R; 728N; 731V; 734Q; 735M; 738I; 739Q; 794E; 797W; 827F; 830E; 831L; 833M; 834N; 835Y; 838E; 841R; 862K; 895M; 899I |
| UBE2D1 | 57Y; 58L; 61K; 65W; 107Y; 108E; 109G; 110R; 111N; 112G; 114W; 115Q; 116Y; 117D; 120T; 137M; 139I; 140A; 144Y; 153Q; 155R; 161R; 162R; 163R; 175K; 178A; 179G; 180L; 181R |
| PANK3 | 17D; 24K; 115G; 116G; 135K; 138E; 189N; 191G; 192S; 193G; 194V; 195S; 207R; 210G; 211T; 253I; 254Y; 258Y; 263L; 267A; 268V; 269A; 299N; 322N; 325R; 336Y; 340Y; 341W |
| AMY2A | 20R; 22V; 23D; 26L; 77D; 81N; 85R; 195R; 233E; 236D; 244S; 245S; 254T; 255E; 256F; 257K; 285G; 286F; 287V; 288P; 295F; 298N; 298S; 300D; 337R; 369E; 370V; 371T; 372I |
| TTR | 13M; 15K; 17L; 18D; 20V; 21R; 24P; 31H; 32V; 33F; 41W; 43P; 44F; 45A; 46S; 52S; 54E; 72E; 78Y; 79W; 82L; 84I; 106T; 108A; 109A; 110L; 115S; 117S; 118T; 119T; 121V |
| MAPK10 | 48K; 49H; 50E; 51M; 52T; 53L; 54K; 55F; 158P; 159K; 160R; 161P; 162T; 163T; 164L; 165N; 166L; 167F; 341V; 342V; 343R; 344P; 345G; 346S; 347L; 348D; 349L; 350P |
| MAOB | 60Y; 84E; 88L; 101G; 102P; 103F; 104P; 119W; 164L; 167L; 168F; 171L; 172C; 198I; 199A; |

| | |
|---|---|
| | 199I; 201T; 206Q; 314T; 316I; 326Y; 327T; 328L; 343F; 398Y; 435Y |
| AKT1 | 53N; 54N; 59Q; 78L; 79Q; 80W; 82T; 83V; 84I; 85E; 205S; 210L; 211T; 264L; 268K; 270V; 271V; 272Y; 273R; 274D; 290I; 291T; 292D; 296C; 326Y |
| UGDH | 131T; 161E; 162F; 163L; 164A; 165E; 220K; 224N; 227L; 231I; 260R; 265F; 266L; 267K; 269S; 272F; 273G; 276C; 277F; 338F; 339K; 442R |
| EPAS1 | 244F; 246S; 248H; 252M; 254F; 277A; 280F; 281Y; 289M; 292S; 293H; 296L; 302V; 304S; 307Y; 309M; 319L; 321T; 323G; 337I; 339C; 341N |
| NT5C2 | 127F; 128I; 129R; 144R; 145D; 147T; 152I; 154N; 292K; 293P; 352H; 354F; 358L; 362K; 424K; 428H; 432M; 436M; 453Q; 456R; 457Y |
| ABL2 | 378R; 382T; 383A; 384V; 386L; 387L; 390A; 475L; 478I; 479A; 480T; 481Y; 508E; 509G; 510C; 511P; 514V; 539F; 542M; 543F; 546S |
| ITK | 356W; 403F; 406E; 407A; 410M; 411M; 413L; 419V; 420Q; 421L; 422Y; 423G; 424V; 435F; 499S; 500D; 501F; 502G; 503M; 505R; 506F |
| CASP7 | 147E; 159I; 160K; 183I; 187R; 211Y; 213I; 214P; 215V; 216E; 221F; 223Y; 227P; 290C; 292V; 294M; 487R; 523Y; 527P; 592V |
| IGF1R | 1033K; 1050E; 1051A; 1054M; 1062V; 1063V; 1065L; 1077V; 1079M; 1133H; 1134R; 1151I; 1152G; 1153D; 1174L; 1189F |
| CASP8 | 395D; 396E; 399F; 400L; 401L; 410V; 411S; 412Y; 413R; 414N; 415P; 419T; 420W; 454D; 455D; 458N; 467T; 468F; 469T |
| PTK2 | 454K; 475M; 483I; 484V; 486L; 534L; 537L; 542F; 544H; 550R; 551N; 552V; 562L; 563G; 564D; 604D; 607M; 608F; 611C |
| HRAS | 11A; 60G; 68R; 72M; 88K; 92D; 93I; 94H; 95Q; 96Y; 97R; 99Q; 102R; 107D; 108D; 109V; 111M; 136S; 137Y; 138G; 139I |
| EIF4E | 45L; 47F; 48F; 49K; 51D; 54K; 58A; 59N; 60L; 61R; 63I; 64S; 75L; 78H; 79I; 80Q; 82S; 83S; 84N; 85L; 86M; 89C; 91Y |
| PANK1 | 416G; 417S; 418G; 419V; 420S; 432R; 435G; 436T; 479Y; 486F; 488L; 492A; 493V; 494A; 524N; 561Y; 565F; 566W |
| PDE10A | 286R; 287C; 288A; 304F; 305D; 324F; 329G; 330I; 331A; 352F; 353N; 356V; 357D; 362Y; 364T; 367I; 383Q; 385V |
| MAP2K2 | 82N; 101K; 119L; 122L; 131V; 132G; 145I; 147M; 193R; 194D; 212D; 213F; 214G; 215V; 216S; 219L; 220I; 223M |
| F7 | 38G; 57H; 60D; 60K; 98T; 99T; 102D; 189D; 190S; 191C; 192K; 195S; 213V; 214S; 215W; 216G; 219G; 220C; 226G |
| FBP1 | 17V; 18M; 20E; 21G; 22R; 24A; 25R; 26G; 27T; 28G; 29E; 30L; 31T; 34L; 112K; 113Y; 140R; 160V; 177M; 178D |

| | |
|---|---|
| KRAS | 7V; 8V; 9V; 10G; 11A; 12C; 13G; 16K; 34P; 58T; 59A; 60G; 61Q; 62E; 63E; 68R; 71Y; 72M; 78F; 96Y; 99Q; 100I |
| PDE4D | 326H; 374S; 439M; 485L; 487N; 495Y; 499T; 502I; 503M; 506F; 534S; 535Q; 538F; 596V; 599F; 600I; 603T |
| MSH2 | 642V; 645Q; 648I; 649A; 650F; 651I; 653N; 670P; 671N; 672M; 673G; 674G; 675K; 676S; 677T; 815Y |
| CHEK1 | 93F; 94D; 95R; 96I; 97E; 98P; 99D; 133P; 173Y; 200A; 204G; 205E; 206L; 207P; 208W; 209D; 217E |
| CDC34 | 18K; 26E; 28F; 45I; 46F; 47G; 48P; 50N; 51T; 52Y; 53Y; 58F; 128I; 131L; 132N; 148Y; 151W; 161Y |
| TYMS | 49D; 50R; 51T; 142F; 160Q; 178I; 179M; 180C; 182W; 183N; 185R; 192L; 196H; 215R; 216S; 258Y |
| PTPN1 | 188P; 189A; 192L; 193N; 196F; 197K; 200E; 276E; 277G; 279K; 280F; 281I; 282M; 291W; 292K |
| ABL1 | 356A; 359L; 360L; 363A; 448L; 451I; 452A; 481E; 482G; 483C; 484P; 487V; 521I; 525V; 529L |
| HBA1 | 35Y; 36F; 37W; 95P; 99K; 100L; 103H; 108N; 137T; 141R; 235Y; 237W; 308N; 495P; 537T; 541R |
| HK1 | 84D; 87G; 88S; 91R; 153T; 155S; 209D; 229I; 231G; 232T; 413D; 414G; 415S; 448G; 449S |
| SERPINC1 | 11K; 12P; 13R; 43A; 44T; 45N; 46R; 47R; 48V; 112S; 113E; 114K; 121F; 122F; 125K; 129R |
| PRKAG1 | 151H; 200T; 203N; 204I; 205A; 225V; 226S; 227A; 229P; 298H; 312I; 314S; 316S; 317D |
| CASP3 | 204Y; 205S; 206W; 207R; 208N; 209S; 210K; 213S; 214W; 248E; 249S; 250F; 251S; 256F |
| ITGAV | 121S; 122Y; 123S; 157V; 180M; 213S; 214R; 215N; 216R; 217D; 218A; 219P; 220E; 253K |
| MALT1 | 344V; 345A; 346L; 379K; 381V; 394A; 397E; 398F; 400L; 401L; 576R; 580W; 715L; 717M |
| GRIA2 | 92I; 104K; 105P; 106F; 107M; 108S; 217S; 218K; 219G; 238V; 239L; 242N; 242S; 247L |
| PRKAA2 | 11V; 18L; 24V; 28G; 29K; 31K; 46I; 48N; 81V; 83R; 88D; 90F; 106T; 107R; 111N; 113V |
| PYGM | 67W; 71Q; 75Y; 282N; 285F; 309R; 310R; 315K; 316F; 317G; 318C; 610A; 612G; 613Y |
| HSPA1A | 15Y; 35N; 37T; 202G; 230G; 268E; 271K; 272R; 275S; 339G; 340S; 342R; 343I; 366D |
| MIF | 32K; 35Q; 36Y; 64I; 65G; 66K; 91P; 95Y; 103A; 108W; 109N; 110N; 111S; 112T; 113F |
| HBB | 35Y; 36F; 37W; 95P; 99K; 100L; 103H; 108N; 137T; 141R; 145H; 225K; 545H; 625K |
| FDPS | 10Y; 57K; 59N; 60R; 63T; 205S; 206F; 239F; 242Q; 246L; 344L; 347K; 348I; 350K |

| | |
|---|---|
| MAPK8 | 178T; 180F; 184P; 197I; 198L; 199G; 230Y; 231I; 234W; 253Q; 255T; 256V; 259Y |
| FBP2 | 17V; 20K; 21G; 24A; 26G; 27T; 28G; 29E; 30L; 31T; 112K; 113Y; 140R; 177T |
| IDH2 | 160L; 164W; 294V; 297V; 298L; 306W; 311Y; 312D; 315V; 316Q; 319I; 320L |
| PPARG | 259E; 280R; 281I; 284G; 285C; 288R; 339V; 341I; 342S; 348M; 353L; 364M |
| PRKAB1 | 18L; 19G; 29K; 31K; 46I; 81V; 83R; 88D; 90F; 106T; 107R; 111N; 113V |
| PDE5A | 612Y; 725L; 765L; 779A; 782V; 783A; 786F; 804L; 816M; 817Q; 820F |
| SIRT3 | 146A; 157F; 173L; 176P; 179I; 180F; 195L; 228Q; 229N; 230I; 231D |
| PRMT3 | 387V; 389D; 392K; 393H; 396R; 420V; 422E; 424L; 466T; 467A; 503L |
| CHRM2 | 80Y; 83Y; 172E; 177Y; 181F; 410N; 414A; 419N; 422W; 423T; 426Y |
| GNPDA1 | 1M; 2K; 151S; 152S; 158R; 159V; 160K; 161T; 184M; 258L; 262H |
| ESR2 | 306L; 309M; 310I; 314K; 324L; 327Q; 328V; 331L; 332E; 335W |
| GLS | 317R; 318F; 320K; 321L; 322F; 323L; 324N; 325E; 327D; 394Y |
| ME2 | 64Q; 67R; 84Y; 88I; 91R; 95L; 127F; 128R; 1127F; 1128R |
| PRPS1 | 99K; 100K; 101D; 102K; 132A; 132S; 135Q; 144N; 146Y |
| ANXA5 | 5L; 118T; 119P; 161R; 203V; 207R; 243S; 244I; 247I |
| CSNK2A1 | 39Y; 40Q; 41L; 67V; 69I; 101V; 103D; 104P; 110A |
| LTA4H | 24R; 25C; 26S; 35T; 36G; 161S; 180D; 182E; 188I |
| AMD1 | 13L; 15E; 111F; 113S; 174D; 176T; 285F; 318Y |
| F2 | 184G; 184Y; 187R; 221D; 221R; 224K; 225Y |
| CYP3A4 | 213F; 214D; 217D; 219F; 220F; 240V |
| MAP2K4 | 3D; 4D; 5E; 6M; 7TPO; 8G; 9Y; 10A |
| CASP1 | 258L; 286R; 331C; 333S; 390E |
| CBS | 439R; 442G; 443F; 444D |
| ADORA2A | 231A |

**Table S3  Parameter settings of grid search**

| Parameter | Lower range | Upper range | Step |
|---|---|---|---|
| n_estimators | 100 | 1000 | 100 |
| eta | 0.025 | 1 | 0.025 |
| max_depth | 1 | 14 | 1 |
| min_child_weight | 1 | 10 | 1 |
| subsample | 0.5 | 1 | 0.05 |
| gamma | 0.5 | 1 | 0.05 |
| colsample_bytree | 0.5 | 1 | 0.05 |

**Table S4 Descriptions of mutation related diseases in allosteric proteins**

| Abbreviation | Disease Name/Description |
|---|---|
| OBESITY | Obesity |
| FPLD3 | Lipodystrophy, familial partial, 3 |
| GLM1 | Glioma 1 |
| IMD12 | Immunodeficiency 12 |
| ECYT4 | Erythrocytosis, familial, 4 |
| PAI-1D | Plasminogen activator inhibitor-1 deficiency |
| CFC3 | Cardiofaciocutaneous syndrome 3 |
| MEL | Melorheostosis, isolated |
| OCNDS | Okur-Chung neurodevelopmental syndrome |
| DEE71 | Developmental and epileptic encephalopathy 71 |
| CASGID | Infantile cataract, skin abnormalities, glutamate excess, and impaired intellectual development |
| GDPAG | Global developmental delay, progressive ataxia, and elevated glutamine |
| MDD | Major depressive disorder |
| CML | Leukemia, chronic myeloid |
| CHDSKM | Congenital heart defects and skeletal malformations syndrome |
| AIS | Androgen insensitivity syndrome |
| SMAX1 | Spinal and bulbar muscular atrophy X-linked 1 |
| PAIS | Androgen insensitivity, partial |
| HYSP1 | Hypospadias 1, X-linked |
| MCLMR | Microcephaly with or without chorioretinopathy, lymphedema, or impaired intellectual development |
| CSTLO | Costello syndrome |
| CMEMS | Congenital myopathy with excess of muscle spindles |
| NMTC2 | Thyroid cancer, non-medullary, 2 |
| BLC | Bladder cancer |
| SFM | Schimmelpenning-Feuerstein-Mims syndrome |

| | |
|---|---|
| AGS5 | Aicardi-Goutieres syndrome 5 |
| CHBL2 | Chilblain lupus 2 |
| D2HGA2 | D-2-hydroxyglutaric aciduria 2 |
| GLM | Glioma |
| LPFS1 | Lymphoproliferative syndrome 1 |
| PRPS1 superactivity | Phosphoribosylpyrophosphate synthetase superactivity |
| CMTX5 | Charcot-Marie-Tooth disease, X-linked recessive, 5 |
| ARTS | ARTS syndrome |
| DFNX1 | Deafness, X-linked, 1 |
| NEDLIB | Neurodevelopmental disorder with language impairment and behavioral abnormalities |
| HEIBAN | Heinz body anemias |
| A-THAL | Alpha-thalassemia |
| HBH | Hemoglobin H disease |
| VDDR3 | Vitamin D-dependent rickets 3 |
| AUTS19 | Autism 19 |
| CPVT4 | Ventricular tachycardia, catecholaminergic polymorphic, 4 |
| LQT14 | Long QT syndrome 14 |
| CFC4 | Cardiofaciocutaneous syndrome 4 |
| AT3D | Antithrombin III deficiency |
| AIS | Androgen insensitivity syndrome |
| SMAX1 | Spinal and bulbar muscular atrophy X-linked 1 |
| PAIS | Androgen insensitivity, partial |
| HYSP1 | Hypospadias 1, X-linked |
| DEE84 | Developmental and epileptic encephalopathy 84 |
| FBP1D | Fructose-1,6-bisphosphatase deficiency |
| HK deficiency | Hexokinase deficiency |
| HMSNR | Neuropathy, hereditary motor and sensory, Russe type |
| RP79 | Retinitis pigmentosa 79 |

| | |
|---|---|
| NEDVIBA | Neurodevelopmental disorder with visual defects and brain anomalies |
| FA2D | Factor II deficiency |
| ISCHSTR | Ischemic stroke (ISCHSTR) |
| THPH1 | Thrombophilia due to thrombin defect |
| RPRGL2 | Pregnancy loss, recurrent, 2 |
| IID | Involvement in disease: A chromosomal aberration involving MAPK10 has been found in a single patient with pharmacoresistant epileptic encephalopathy. Translocation t(Y;4)(q11.2;q21) which causes MAPK10 truncation |
| MODY2 | Maturity-onset diabetes of the young 2 |
| HHF3 | Hyperinsulinemic hypoglycemia, familial, 3 |
| T2D | Type 2 diabetes mellitus |
| PNDM1 | Diabetes mellitus, permanent neonatal, 1 |
| ODG8 | Ovarian dysgenesis 8 |
| IGF1RES | Insulin-like growth factor 1 resistance |
| IID2 | Involvement in disease: Aberrant PTK2/FAK1 expression may play a role in cancer cell proliferation, migration and invasion, in tumor formation and metastasis. PTK2/FAK1 overexpression is seen in many types of cancer |
| IID3 | Involvement in disease: Genetic variations in PDE4D might be associated with susceptibility to stroke. |
| ACRDYS2 | Acrodysostosis 2, with or without hormone resistance |
| FA7D | Factor VII deficiency |
| AML | Leukemia, acute myelogenous |
| JMML | Leukemia, juvenile myelomonocytic |
| NS3 | Noonan syndrome 3 |
| GASC | Gastric cancer |
| CFC2 | Cardiofaciocutaneous syndrome 2 |
| OES | Oculoectodermal syndrome |
| SFM | Schimmelpenning-Feuerstein-Mims syndrome |
| AMYL-TTR | Amyloidosis, transthyretin-related |
| DTTRH | Hyperthyroxinemia, dystransthyretinemic |

| | |
|---|---|
| CTS1 | Carpal tunnel syndrome 1 |
| CRC | Colorectal cancer |
| LNCR | Lung cancer |
| NHL | Familial non-Hodgkin lymphoma |
| CFC1 | Cardiofaciocutaneous syndrome 1 |
| NS7 | Noonan syndrome 7 |
| LPRD3 | LEOPARD syndrome 3 |
| CASP8D | Caspase-8 deficiency |
| HEIBAN | Heinz body anemias |
| B-THAL | Beta-thalassemia |
| SKCA | Sickle cell disease |
| B-THALIB | Beta-thalassemia, dominant, inclusion body type |
| GSD5 | Glycogen storage disease 5 |
| RASJ | Rheumatoid arthritis systemic juvenile |
| CBSD | Cystathionine beta-synthase deficiency |
| IID4 | Involvement in disease: in certain aggressive cases of activated B cell-like diffuse large B-cell lymphoma (ABC-DLBCL), plays a role in the cytoplasmic sequestration of misfolded N-terminal mutated PRDM1 proteins, promotes their association with SYNV1/HRD1 and degradation through the SYNV1-proteasome pathway. HSPA1A inhibition restores PRDM1 nuclear localization and transcriptional activity in lymphoma cell lines and suppresses growth in xenografts |
| BC | Breast cancer |
| CRC | Colorectal cancer |
| PROTEUSS | Proteus syndrome |
| CWS6 | Cowden syndrome 6 |
| FDAH | Hyperthyroxinemia, familial dysalbuminemic |
| ANALBA | Analbuminemia |
| SPG45 | Spastic paraplegia 45, autosomal recessive |
| GSD6 | Glycogen storage disease 6 |
| LYNCH1 | Lynch syndrome 1 |

| | |
|---|---|
| MRTES | Muir-Torre syndrome |
| ENDMC | Endometrial cancer |
| MMRCS2 | Mismatch repair cancer syndrome 2 |
| CRC | Colorectal cancer |
| POROK9 | Porokeratosis 9, multiple types |
| IOLOD | Dyskinesia, limb and orofacial, infantile-onset |
| ADSD2 | Striatal degeneration, autosomal dominant 2 |
| CORLK | Leukodystrophy, childhood-onset, remitting |
| PKHYP | Pyruvate kinase hyperactivity |
| PKRD | Pyruvate kinase deficiency of red cells |
| RPRGL3 | Pregnancy loss, recurrent, 3 |
| DKCD | Dyskeratosis congenita, digenic |

**Table S5 The maximum, minimum, median and average output scores of different prediction methods**

| Method | Maximum | Minimum | Median | Mean |
|---|---|---|---|---|
| MetaMutPre | 1.219 | -0.0791 | 0.410 | 0.464 |
| AlphaMissence | 1.000 | 0.000 | 0.450 | 0.514 |
| AlphaScore | 0.666 | 0.885 | 0.791 | 0.796 |
| SPRI | 0.996 | 0.000 | 0.734 | 0.672 |
| ESM1b | 16.896 | -30.945 | -7.355 | -8.183 |
| gMVP | 1.000 | 0.000 | 0.500 | 0.500 |
| DEOGEN2 | 0.999 | 0.000 | 0.152 | 0.245 |
| REVEL | 1.000 | 0.000 | 0.217 | 0.302 |
| CADD | 21.370 | -18.155 | 2.848 | 2.642 |

**Table S6  Optimal threshold determined by Youden index.**

| Method | Threshold1 | Threshold2 | Threshold3 |
|---|---|---|---|
| MetaXGBoost | 0.5526 | 0.6377 | 0.9412 |
| gMVP | 0.812854 | 0.812854 | 0.9546 |
| REVEL | 0.695 | 0.751 | 0.85 |
| AlphaMissence | 0.7334 | 0.7334 | 0.9865 |
| CADD | 3.25832 | 3.25832 | 4.071655 |
| ESM1b | -9.369 | -9.661 | -8.711 |
| SPRI | 0.652 | 0.652 | 0.538 |
| DEOGEN2 | 0.827048 | 0.836823 | 0.816905 |

Note: Threshold1, Threshold2 and Threshold3 are the optimal thresholds for whole gene regions, allosteric-protein-encoded gene regions and allosteric sites, respectively.

**Table S7 Potential pathogenic mutation on allosteric sites predicted by MetaMutPre**

| Gene | Mutation | Clinvar | Gene | Mutation | Clinvar |
|---|---|---|---|---|---|
| ABL1 | P484R | - | ABL1 | P484S | - |
| ABL1 | P484T | - | ABL2 | L386P | - |
| ABL2 | L387H | - | ABL2 | L387P | - |
| ABL2 | L387R | - | ABL2 | A390D | - |
| ABL2 | A390P | - | ABL2 | L475S | - |
| ABL2 | L475W | - | ABL2 | I478N | - |
| ABL2 | I478S | - | ABL2 | A479D | - |
| ABL2 | A479P | - | ABL2 | T480I | - |
| ABL2 | T480N | - | ABL2 | T480P | - |
| ABL2 | C510F | - | ABL2 | C510G | - |
| ABL2 | C510R | - | ABL2 | C510S | - |
| ABL2 | C510Y | - | ABL2 | V514D | - |
| ABL2 | V514G | - | AKT1 | L78P | - |
| AKT1 | Q79P | - | AKT1 | W80C | - |
| AKT1 | W80G | - | AKT1 | W80R | - |
| AKT1 | W80S | - | AKT1 | I84N | - |
| AKT1 | I84S | - | AKT1 | E85A | - |
| AKT1 | E85G | - | AKT1 | E85K | - |
| AKT1 | E85Q | - | AKT1 | E85V | - |
| AKT1 | L210P | - | AKT1 | L264P | - |
| AKT1 | L264Q | - | AKT1 | L264R | - |
| AKT1 | V270E | - | AKT1 | V270G | - |
| AKT1 | V271E | - | AKT1 | V271G | - |
| AKT1 | Y272D | - | AKT1 | R273G | - |
| AKT1 | R273Q | - | AKT1 | D274A | - |
| AKT1 | D274G | - | AKT1 | D274H | - |
| AKT1 | D274N | - | AKT1 | D274V | - |

| | | | | | |
|---|---|---|---|---|---|
| AKT1 | D274Y | - | AKT1 | I290S | - |
| AKT1 | T291K | - | AKT1 | D292A | - |
| AKT1 | D292G | - | AKT1 | D292H | - |
| AKT1 | D292N | - | AKT1 | D292V | - |
| AKT1 | D292Y | - | AKT1 | C296F | - |
| AKT1 | C296R | - | AKT1 | C296Y | - |
| AMY2A | W295G | - | AMY2A | W295R | - |
| AR | L713F | * | AR | L713H | - |
| AR | L713P | - | AR | L713R | - |
| AR | V717A | - | AR | V717D | - |
| AR | V717G | - | AR | K718E | - |
| AR | K721E | - | AR | K721Q | - |
| AR | K721T | - | AR | P724A | - |
| AR | P724H | - | AR | P724L | * |
| AR | P724R | - | AR | P724S | * |
| AR | P724T | - | AR | G725C | - |
| AR | G725D | - | AR | G725R | - |
| AR | G725S | - | AR | G725V | - |
| AR | F726C | - | AR | F726S | - |
| AR | F726V | - | AR | R727G | - |
| AR | R727S | - | AR | V731E | - |
| AR | V731G | - | AR | Q734E | - |
| AR | Q734K | - | AR | Q734L | - |
| AR | Q734P | - | AR | Q734R | - |
| AR | M735I | - | AR | M735K | - |
| AR | M735R | - | AR | M735T | - |
| AR | M735V | - | AR | I738F | - |
| AR | I738N | - | AR | I738S | - |
| AR | I738T | * | AR | Q739E | - |
| AR | Q739K | - | AR | Q739P | - |
| AR | L831H | - | AR | L831P | - |
| AR | L831R | - | AR | Y835C | * |
| AR | Y835D | - | AR | Y835H | - |
| AR | Y835N | - | AR | Y835S | - |
| AR | E838K | - | AR | E838Q | - |
| AR | E838V | - | AR | M895I | - |
| AR | M895K | - | AR | M895R | - |
| AR | M895T | - | AR | M895V | - |
| AR | I899F | - | AR | I899N | - |
| AR | I899S | - | BRAF | I463N | - |
| BRAF | I463S | * | BRAF | S467L | - |

| | | | | | |
|---|---|---|---|---|---|
| BRAF | S467P | - | BRAF | T470P | * |
| BRAF | V471A | - | BRAF | V471D | - |
| BRAF | V471F | * | BRAF | V471G | - |
| BRAF | Y472D | - | BRAF | Y472N | - |
| BRAF | Y472S | - | BRAF | V480E | - |
| BRAF | V480G | - | BRAF | A481E | * |
| BRAF | A481G | - | BRAF | A481P | - |
| BRAF | A481T | - | BRAF | A481V | - |
| BRAF | V482E | - | BRAF | V482G | - |
| BRAF | K483E | * | BRAF | K483I | - |
| BRAF | K483Q | * | BRAF | K483R | - |
| BRAF | K483T | * | BRAF | A497P | - |
| BRAF | E501A | * | BRAF | E501G | * |
| BRAF | E501K | * | BRAF | E501Q | * |
| BRAF | E501V | * | BRAF | V504E | - |
| BRAF | V504G | - | BRAF | L505H | - |
| BRAF | L505P | - | BRAF | L505R | - |
| BRAF | K507E | - | BRAF | N512D | - |
| BRAF | N512H | - | BRAF | N512I | - |
| BRAF | N512S | - | BRAF | N512T | - |
| BRAF | N512Y | - | BRAF | I513F | - |
| BRAF | I513N | - | BRAF | I513S | - |
| BRAF | I513T | - | BRAF | L514P | - |
| BRAF | L514Q | - | BRAF | L514R | - |
| BRAF | L515P | - | BRAF | L515R | - |
| BRAF | F516C | - | BRAF | F516I | - |
| BRAF | F516S | - | BRAF | A526D | - |
| BRAF | A526P | - | BRAF | I527N | - |
| BRAF | I527S | - | BRAF | I527T | - |
| BRAF | V528D | - | BRAF | V528F | - |
| BRAF | V528G | - | BRAF | T529I | - |
| BRAF | T529N | - | BRAF | T529P | - |
| BRAF | Q530K | - | BRAF | Q530P | - |
| BRAF | W531C | * | BRAF | W531G | - |
| BRAF | W531L | * | BRAF | W531R | - |
| BRAF | W531S | * | BRAF | C532F | - |
| BRAF | C532G | - | BRAF | C532R | - |
| BRAF | C532S | - | BRAF | C532Y | * |
| BRAF | E533G | - | BRAF | E533K | - |
| BRAF | E533V | - | BRAF | S535F | - |
| BRAF | S535P | - | BRAF | S535Y | - |

| | | | | | |
|---|---|---|---|---|---|
| BRAF | S536C | - | BRAF | S536G | - |
| BRAF | S536I | - | BRAF | S536N | - |
| BRAF | Y566C | - | BRAF | Y566D | - |
| BRAF | Y566H | - | BRAF | Y566N | - |
| BRAF | Y566S | - | BRAF | L567F | - |
| BRAF | L567S | - | BRAF | S571P | - |
| BRAF | I572F | - | BRAF | I572N | - |
| BRAF | I572S | - | BRAF | I572T | - |
| BRAF | I573F | - | BRAF | I573N | - |
| BRAF | I573S | - | BRAF | I573T | - |
| BRAF | H574D | * | BRAF | H574L | - |
| BRAF | H574N | - | BRAF | H574P | - |
| BRAF | H574R | - | BRAF | H574Y | * |
| BRAF | N581D | * | BRAF | N581H | * |
| BRAF | N581I | - | BRAF | N581S | * |
| BRAF | N581T | * | BRAF | N581Y | - |
| BRAF | I582K | - | BRAF | I582R | - |
| BRAF | I582T | - | BRAF | F583C | - |
| BRAF | F583S | - | BRAF | F583V | - |
| BRAF | K591E | - | BRAF | K591I | - |
| BRAF | K591Q | - | BRAF | K591R | - |
| BRAF | K591T | - | BRAF | I592K | - |
| BRAF | I592R | - | BRAF | I592T | - |
| BRAF | G593C | - | BRAF | G593D | - |
| BRAF | G593R | - | BRAF | G593V | - |
| BRAF | D594A | * | BRAF | D594G | * |
| BRAF | D594H | * | BRAF | D594N | * |
| BRAF | D594V | * | BRAF | D594Y | - |
| BRAF | F595C | - | BRAF | F595I | - |
| BRAF | F595S | * | BRAF | F595V | - |
| BRAF | F595Y | - | BRAF | G596A | - |
| BRAF | G596C | * | BRAF | G596D | * |
| BRAF | G596R | * | BRAF | G596S | * |
| BRAF | G596V | * | BRAF | L597P | - |
| BRAF | L597Q | * | BRAF | L597R | * |
| BRAF | A598D | - | BRAF | A598G | - |
| BRAF | A598P | - | BRAF | A598T | - |
| BRAF | A598V | - | BRAF | T599P | - |
| BRAF | K601E | * | CDK2 | K33E | - |
| CDK2 | K33Q | - | CDK2 | K33T | - |
| CDK2 | V64D | - | CDK2 | L66P | - |

| | | | | | |
|---|---|---|---|---|---|
| CDK2 | L78P | - | CDK2 | A144E | - |
| CDK2 | D145A | - | CDK2 | D145G | - |
| CDK2 | D145H | - | CDK2 | D145V | - |
| CDK2 | D145Y | - | CDK2 | F146S | - |
| CDK2 | L148P | - | CDK2 | L148Q | - |
| CDK2 | L148R | - | CHKA | L124P | - |
| CHKA | R146G | - | CHKA | R146P | - |
| CHKA | E180A | - | CHKA | E180G | - |
| CHKA | E180K | - | CHKA | E180Q | - |
| CHKA | E180V | - | CHKA | P194L | - |
| CHKA | P194Q | - | CHKA | P194R | - |
| CHKA | P194S | * | CHKA | E206A | - |
| CHKA | E206G | - | CHKA | E206K | - |
| CHKA | E206V | - | CHKA | L214S | - |
| CHKA | D306A | - | CHKA | D306G | - |
| CHKA | D306H | - | CHKA | D306N | - |
| CHKA | D306V | - | CHKA | D306Y | - |
| CHKA | Q308K | - | CHKA | G310C | - |
| CHKA | G310R | - | CHKA | I329N | - |
| CHKA | I329S | - | CHKA | D330A | - |
| CHKA | D330G | - | CHKA | D330H | - |
| CHKA | D330N | - | CHKA | D330V | - |
| CHKA | D330Y | - | CHKA | E349A | - |
| CHKA | E349G | - | CHKA | E349K | - |
| CHKA | E349Q | - | CHKA | E349V | - |
| CHKA | P370R | - | CHKA | W420C | - |
| CHKA | W420L | - | CHKA | W420R | - |
| CHKA | W420S | - | CHKA | W423G | - |
| CHKA | W423S | - | CHRM2 | W422C | - |
| CHRM2 | W422R | - | CHRM2 | W422S | - |
| CSNK2A1 | Y39D | - | CSNK2A1 | Q40P | - |
| CSNK2A1 | L41P | - | CSNK2A1 | L41Q | - |
| CSNK2A1 | L41R | - | CSNK2A1 | V67D | - |
| CSNK2A1 | I69N | - | CSNK2A1 | V101E | - |
| CSNK2A1 | D103A | - | CSNK2A1 | D103G | - |
| CSNK2A1 | A110D | - | CSNK2A1 | A110P | - |
| EPAS1 | Y307D | - | EPAS1 | M309R | - |
| ESR2 | L306S | - | ESR2 | L306W | - |
| ESR2 | M309K | - | ESR2 | M309R | - |
| ESR2 | I310N | - | ESR2 | I310S | - |
| ESR2 | K314E | - | ESR2 | Q327E | - |

| | | | | | |
|---|---|---|---|---|---|
| ESR2 | Q327K | - | ESR2 | Q327P | - |
| ESR2 | L331S | - | ESR2 | L331W | - |
| ESR2 | W335C | - | ESR2 | W335G | - |
| ESR2 | W335L | - | ESR2 | W335R | - |
| ESR2 | W335S | - | F2 | Y221C | - |
| F2 | Y221D | - | F2 | Y221S | - |
| F7 | C195F | - | F7 | C195G | - |
| F7 | C195R | * | F7 | C195S | - |
| F7 | C195Y | - | F7 | V214E | - |
| F7 | G215V | - | F7 | C219R | - |
| F7 | W226C | - | F7 | W226G | - |
| F7 | W226L | - | F7 | W226R | - |
| F7 | W226S | - | FBP1 | V18D | - |
| FBP1 | V18G | - | FBP1 | L31S | - |
| FBP1 | L34P | - | FBP1 | L34Q | - |
| FBP1 | L34R | - | FBP1 | G112C | - |
| FBP1 | G112D | - | FBP1 | G112R | - |
| FBP1 | G112V | - | FBP1 | Y140D | - |
| FBP1 | Y140H | - | FBP1 | Y140N | - |
| FBP1 | Y140S | - | FBP1 | L160P | - |
| FBP1 | L160R | - | FBP1 | A177P | - |
| FBP2 | L31P | - | FBP2 | L31R | - |
| FBP2 | G112V | - | FBP2 | Y140D | - |
| FBP2 | Y140N | - | FBP2 | Y140S | - |
| FBP2 | S177F | - | FBP2 | S177P | - |
| GCK | Y215D | - | GCK | Y215N | - |
| GCK | Y215S | - | GLS | F318C | - |
| GLS | F318I | - | GLS | F318S | - |
| GLS | F318V | - | GLS | L321P | - |
| GLS | L321Q | - | GLS | L321R | - |
| GLS | L323S | - | GLS | Y394D | - |
| GLS | Y394N | - | GRIA2 | Y92D | - |
| GRIA2 | Y92N | - | GRIA2 | C104F | - |
| GRIA2 | C104R | - | GRIA2 | C104Y | - |
| GRIA2 | L107P | - | GRIA2 | L107R | - |
| GRIA2 | I219N | - | GRIA2 | I219S | - |
| GRIA2 | A238E | - | GRIA2 | A238P | - |
| GRIA2 | F242S | - | HBA1 | D95A | * |
| HBA1 | D95G | - | HBA1 | D95H | - |
| HBA1 | D95V | - | HBA1 | D95Y | * |
| HBA1 | V108E | - | HBA1 | L137P | * |

| | | | | | |
|---|---|---|---|---|---|
| HBA1 | L137Q | - | HBA1 | L137R | * |
| HBB | V99A | - | HBB | V99E | - |
| HBB | V99G | * | HBB | D100A | * |
| HBB | D100G | * | HBB | D100H | * |
| HBB | D100V | * | HBB | D100Y | * |
| HBB | N103D | - | HBB | G108D | - |
| HBB | A141D | * | HBB | A141P | - |
| HRAS | G60A | - | HRAS | G60C | - |
| HRAS | G60D | * | HRAS | G60R | - |
| HRAS | G60S | * | HRAS | G60V | * |
| HRAS | R68G | - | HRAS | R68P | - |
| HRAS | M72K | - | HRAS | M72R | - |
| HRAS | M72T | - | HRAS | I93N | - |
| HRAS | I93S | - | HRAS | Y96D | - |
| HRAS | Y96N | - | HRAS | Y96S | - |
| HRAS | R97G | - | HRAS | R97T | - |
| HRAS | Q99P | - | HRAS | V109E | - |
| HRAS | V109G | - | HRAS | M111R | - |
| IDH2 | W164C | - | IDH2 | W164G | - |
| IDH2 | W164L | - | IDH2 | W164R | - |
| IDH2 | W164S | - | IDH2 | V294E | - |
| IDH2 | V294G | - | IDH2 | L298H | - |
| IDH2 | L298P | - | IDH2 | L298R | - |
| IDH2 | W306C | - | IDH2 | W306G | - |
| IDH2 | W306L | - | IDH2 | W306R | - |
| IDH2 | W306S | - | IDH2 | Y311C | - |
| IDH2 | Y311D | - | IDH2 | Y311H | - |
| IDH2 | Y311N | - | IDH2 | Y311S | - |
| IDH2 | D312G | - | IDH2 | D312V | - |
| IDH2 | V315A | - | IDH2 | V315E | - |
| IDH2 | V315G | - | IDH2 | Q316E | - |
| IDH2 | Q316K | - | IDH2 | Q316P | - |
| IDH2 | Q316R | - | IDH2 | L320P | - |
| IDH2 | L320Q | - | IDH2 | L320R | - |
| IGF1R | K1033E | - | IGF1R | K1033I | - |
| IGF1R | K1033Q | - | IGF1R | K1033R | - |
| IGF1R | K1033T | - | IGF1R | E1050A | - |
| IGF1R | E1050G | - | IGF1R | E1050K | - |
| IGF1R | E1050Q | - | IGF1R | E1050V | - |
| IGF1R | A1051D | - | IGF1R | A1051P | - |
| IGF1R | M1054I | * | IGF1R | M1054K | - |

| | | | | | |
|---|---|---|---|---|---|
| IGF1R | M1054R | - | IGF1R | M1054T | - |
| IGF1R | M1054V | - | IGF1R | V1062A | - |
| IGF1R | V1062E | - | IGF1R | V1062G | - |
| IGF1R | V1063A | - | IGF1R | V1063E | - |
| IGF1R | V1063G | - | IGF1R | L1065S | - |
| IGF1R | L1065W | - | IGF1R | V1077D | - |
| IGF1R | V1077G | - | IGF1R | M1079K | - |
| IGF1R | M1079R | - | IGF1R | M1079T | - |
| IGF1R | H1133D | - | IGF1R | H1133L | - |
| IGF1R | H1133N | - | IGF1R | H1133P | - |
| IGF1R | H1133R | - | IGF1R | H1133Y | - |
| IGF1R | R1134G | - | IGF1R | R1134I | - |
| IGF1R | R1134K | - | IGF1R | R1134S | - |
| IGF1R | R1134T | - | IGF1R | I1151F | - |
| IGF1R | I1151N | - | IGF1R | I1151S | - |
| IGF1R | I1151T | - | IGF1R | G1152E | - |
| IGF1R | G1152R | * | IGF1R | G1152V | - |
| IGF1R | D1153A | - | IGF1R | D1153G | - |
| IGF1R | D1153H | - | IGF1R | D1153N | - |
| IGF1R | D1153V | - | IGF1R | D1153Y | - |
| IGF1R | L1174P | - | IGF1R | L1174Q | - |
| IGF1R | L1174R | - | IGF1R | F1189C | - |
| IGF1R | F1189I | - | IGF1R | F1189S | - |
| IGF1R | F1189V | - | ITGAL | L157P | - |
| ITGAL | L157Q | - | ITGAL | L157R | - |
| ITGAL | T231I | - | ITGAL | T231P | - |
| ITGAL | T233A | - | ITGAL | T233I | - |
| ITGAL | T233P | - | ITGAL | V258E | - |
| ITGAL | L259P | - | ITGAL | L259R | - |
| ITGAL | S304P | - | ITK | W356C | - |
| ITK | W356G | - | ITK | F403S | - |
| ITK | E406A | - | ITK | E406G | - |
| ITK | E406K | - | ITK | E406Q | - |
| ITK | E406V | - | ITK | M410K | - |
| ITK | M410R | - | ITK | L413P | - |
| ITK | L413R | - | ITK | V419E | - |
| ITK | V419G | - | ITK | L421P | - |
| ITK | L421Q | - | ITK | L421R | - |
| ITK | G423R | - | ITK | V424E | - |
| ITK | S499F | - | ITK | S499P | - |
| ITK | S499Y | - | ITK | D500A | - |

| | | | | | |
|---|---|---|---|---|---|
| ITK | D500G | - | ITK | D500H | - |
| ITK | D500V | - | ITK | D500Y | - |
| ITK | F501C | - | ITK | F501I | - |
| ITK | F501S | - | ITK | F501V | - |
| ITK | G502A | - | ITK | G502E | - |
| ITK | G502R | - | ITK | G502V | - |
| ITK | G502W | - | ITK | M503K | - |
| ITK | M503R | - | ITK | M503T | - |
| ITK | R505G | - | ITK | R505M | - |
| ITK | R505T | - | ITK | R505W | - |
| KIF11 | Y104C | - | KIF11 | Y104D | - |
| KIF11 | Y104H | - | KIF11 | Y104N | - |
| KIF11 | Y104S | - | KIF11 | T112A | - |
| KIF11 | T112I | - | KIF11 | T112N | - |
| KIF11 | T112P | - | KIF11 | G117A | - |
| KIF11 | G117C | - | KIF11 | G117D | - |
| KIF11 | G117R | - | KIF11 | G117S | - |
| KIF11 | G117V | - | KIF11 | W127C | - |
| KIF11 | D130N | - | KIF11 | I136F | - |
| KIF11 | I136N | - | KIF11 | I136S | - |
| KIF11 | P137L | - | KIF11 | P137Q | - |
| KIF11 | P137R | - | KIF11 | P137S | - |
| KIF11 | P137T | - | KIF11 | L160P | - |
| KIF11 | L160Q | - | KIF11 | L160R | - |
| KIF11 | L171H | - | KIF11 | L171P | - |
| KIF11 | L171R | - | KIF11 | L172H | - |
| KIF11 | L172P | - | KIF11 | L172R | - |
| KIF11 | V210D | - | KIF11 | L214S | - |
| KIF11 | G217E | - | KIF11 | G217R | - |
| KIF11 | G217V | - | KIF11 | G217W | - |
| KIF11 | R221G | - | KIF11 | A224P | - |
| KIF11 | M228K | - | KIF11 | M228R | - |
| KIF11 | N229D | - | KIF11 | N229H | - |
| KIF11 | N229I | - | KIF11 | N229T | - |
| KIF11 | N229Y | - | KIF11 | S232A | - |
| KIF11 | S232C | - | KIF11 | S232F | - |
| KIF11 | S232P | - | KIF11 | S232T | - |
| KIF11 | S232Y | - | KIF11 | F239C | - |
| KIF11 | F239I | - | KIF11 | F239S | - |
| KIF11 | F239V | - | KIF11 | L266F | - |
| KIF11 | L266H | - | KIF11 | L266P | - |

| | | | | | |
|---|---|---|---|---|---|
| KIF11 | L266R | - | KIF11 | S269C | - |
| KIF11 | S269G | - | KIF11 | S269I | - |
| KIF11 | S269N | - | KIF11 | S269T | - |
| KIF11 | E270A | - | KIF11 | E270G | - |
| KIF11 | E270K | - | KIF11 | E270Q | - |
| KIF11 | E270V | - | KIF11 | I288K | - |
| KIF11 | I288R | - | KIF11 | I288T | - |
| KIF11 | N289D | - | KIF11 | N289H | - |
| KIF11 | N289I | - | KIF11 | N289S | - |
| KIF11 | N289T | - | KIF11 | N289Y | - |
| KIF11 | L292P | - | KIF11 | L292Q | - |
| KIF11 | L292R | - | KIF11 | L293S | - |
| KIF11 | L293W | - | KIF11 | L295F | - |
| KIF11 | L295S | - | KIF11 | L295W | - |
| KIF11 | G296A | - | KIF11 | G296E | - |
| KIF11 | G296R | - | KIF11 | G296V | - |
| KIF11 | R297G | - | KIF11 | I299F | - |
| KIF11 | I299N | - | KIF11 | I299S | - |
| KIF11 | I299T | - | KIF11 | I332K | - |
| KIF11 | I332R | - | KIF11 | I332T | - |
| KIF11 | Y352C | - | KIF11 | Y352D | - |
| KIF11 | Y352H | - | KIF11 | Y352N | - |
| KIF11 | Y352S | - | KIF11 | A353D | - |
| KIF11 | A353P | - | KIF11 | A353V | - |
| KIF11 | A356E | - | KIF11 | A356P | - |
| KRAS | V7E | - | KRAS | V7G | - |
| KRAS | V8A | - | KRAS | V8E | - |
| KRAS | V8G | - | KRAS | V9A | - |
| KRAS | V9D | - | KRAS | V9G | - |
| KRAS | G10A | - | KRAS | G10E | - |
| KRAS | G10R | - | KRAS | G10V | - |
| KRAS | G12A | * | KRAS | G12C | * |
| KRAS | G12D | * | KRAS | G12R | * |
| KRAS | G12S | * | KRAS | G12V | * |
| KRAS | G13C | * | KRAS | G13R | * |
| KRAS | G13V | * | KRAS | K16E | - |
| KRAS | K16Q | - | KRAS | K16R | - |
| KRAS | K16T | - | KRAS | P34L | * |
| KRAS | P34Q | * | KRAS | P34R | * |
| KRAS | P34T | - | KRAS | T58A | - |
| KRAS | T58I | * | KRAS | T58K | - |

| | | | | | |
|---|---|---|---|---|---|
| KRAS | T58P | - | KRAS | T58R | - |
| KRAS | A59E | - | KRAS | A59P | - |
| KRAS | G60A | - | KRAS | G60C | - |
| KRAS | G60D | - | KRAS | G60R | * |
| KRAS | G60S | * | KRAS | G60V | * |
| KRAS | Q61E | * | KRAS | Q61K | * |
| KRAS | Q61L | * | KRAS | Q61P | * |
| KRAS | R68G | - | KRAS | Y71C | - |
| KRAS | Y71D | * | KRAS | Y71H | * |
| KRAS | Y71N | - | KRAS | Y71S | - |
| KRAS | M72K | - | KRAS | M72R | - |
| KRAS | M72T | - | KRAS | F78C | - |
| KRAS | F78S | - | KRAS | Y96D | - |
| KRAS | Y96N | - | KRAS | Y96S | - |
| KRAS | Q99P | - | KRAS | I100F | - |
| KRAS | I100N | - | KRAS | I100S | - |
| KRAS | I100T | - | MALT1 | V344D | - |
| MALT1 | V344G | - | MALT1 | A345D | - |
| MALT1 | A345P | - | MALT1 | L346H | - |
| MALT1 | L346P | - | MALT1 | L346R | - |
| MALT1 | A394P | - | MALT1 | W580C | - |
| MALT1 | W580R | - | MALT1 | W580S | * |
| MAOB | Y60C | - | MAOB | Y60D | - |
| MAOB | Y60F | - | MAOB | Y60H | - |
| MAOB | Y60N | - | MAOB | Y60S | - |
| MAOB | L88P | - | MAOB | W119S | - |
| MAOB | L167P | - | MAOB | F168S | - |
| MAOB | I198N | - | MAOB | I198S | - |
| MAOB | Q206E | - | MAOB | Q206K | - |
| MAOB | Q206L | - | MAOB | Q206P | - |
| MAOB | T327K | - | MAOB | T327P | - |
| MAOB | T327R | - | MAOB | L328S | - |
| MAOB | F343S | - | MAOB | Y435C | - |
| MAOB | Y435D | - | MAOB | Y435H | - |
| MAOB | Y435N | - | MAOB | Y435S | - |
| MAP2K1 | G77A | - | MAP2K1 | G77C | - |
| MAP2K1 | G77D | - | MAP2K1 | G77R | - |
| MAP2K1 | G77S | - | MAP2K1 | G77V | - |
| MAP2K1 | G79D | - | MAP2K1 | G79R | - |
| MAP2K1 | G79V | - | MAP2K1 | G80C | - |
| MAP2K1 | G80D | - | MAP2K1 | G80R | - |

| | | | | | |
|---|---|---|---|---|---|
| MAP2K1 | G80V | - | MAP2K1 | K97E | - |
| MAP2K1 | K97N | - | MAP2K1 | K97Q | - |
| MAP2K1 | K97R | - | MAP2K1 | K97T | - |
| MAP2K1 | I99N | - | MAP2K1 | I99S | - |
| MAP2K1 | L115P | - | MAP2K1 | L115Q | - |
| MAP2K1 | L115R | - | MAP2K1 | L118P | - |
| MAP2K1 | L118Q | - | MAP2K1 | L118R | - |
| MAP2K1 | V127A | - | MAP2K1 | V127E | - |
| MAP2K1 | V127G | - | MAP2K1 | Y130D | - |
| MAP2K1 | Y130N | * | MAP2K1 | Y130S | - |
| MAP2K1 | I141N | - | MAP2K1 | I141S | - |
| MAP2K1 | I141T | - | MAP2K1 | C142R | - |
| MAP2K1 | M143I | - | MAP2K1 | M143K | - |
| MAP2K1 | M143R | - | MAP2K1 | M143T | - |
| MAP2K1 | M143V | * | MAP2K1 | E144A | - |
| MAP2K1 | E144G | - | MAP2K1 | E144K | - |
| MAP2K1 | E144Q | - | MAP2K1 | E144V | - |
| MAP2K1 | H188D | - | MAP2K1 | H188L | - |
| MAP2K1 | H188N | - | MAP2K1 | H188P | - |
| MAP2K1 | H188R | - | MAP2K1 | H188Y | - |
| MAP2K1 | R189G | - | MAP2K1 | R189I | - |
| MAP2K1 | R189K | - | MAP2K1 | R189S | - |
| MAP2K1 | R189T | - | MAP2K1 | D190A | - |
| MAP2K1 | D190G | - | MAP2K1 | D190H | - |
| MAP2K1 | D190N | - | MAP2K1 | D190V | - |
| MAP2K1 | D190Y | - | MAP2K1 | V191A | - |
| MAP2K1 | V191D | - | MAP2K1 | V191G | - |
| MAP2K1 | K192E | - | MAP2K1 | K192Q | - |
| MAP2K1 | K192R | - | MAP2K1 | K192T | - |
| MAP2K1 | N195D | - | MAP2K1 | N195H | - |
| MAP2K1 | N195I | - | MAP2K1 | N195S | - |
| MAP2K1 | N195T | - | MAP2K1 | N195Y | - |
| MAP2K1 | C207F | - | MAP2K1 | C207G | - |
| MAP2K1 | C207R | - | MAP2K1 | C207S | - |
| MAP2K1 | C207Y | - | MAP2K1 | D208A | - |
| MAP2K1 | D208G | - | MAP2K1 | D208H | - |
| MAP2K1 | D208N | - | MAP2K1 | D208V | - |
| MAP2K1 | D208Y | - | MAP2K1 | F209C | - |
| MAP2K1 | F209I | - | MAP2K1 | F209S | - |
| MAP2K1 | F209V | - | MAP2K1 | G210A | - |
| MAP2K1 | G210E | - | MAP2K1 | G210R | - |

| | | | | | |
|---|---|---|---|---|---|
| MAP2K1 | G210V | - | MAP2K1 | G210W | - |
| MAP2K1 | V211D | * | MAP2K1 | V211G | - |
| MAP2K1 | S212C | - | MAP2K1 | S212G | - |
| MAP2K1 | S212I | - | MAP2K1 | S212N | - |
| MAP2K1 | S212T | - | MAP2K1 | G213E | - |
| MAP2K1 | G213V | - | MAP2K1 | G213W | - |
| MAP2K1 | L215H | - | MAP2K1 | L215P | * |
| MAP2K1 | L215R | - | MAP2K1 | D217V | - |
| MAP2K1 | S222F | - | MAP2K1 | S222P | - |
| MAP2K1 | S222Y | - | MAP2K1 | F223C | - |
| MAP2K1 | F223I | - | MAP2K1 | F223S | - |
| MAP2K1 | F223V | - | MAP2K1 | G225A | - |
| MAP2K1 | G225C | - | MAP2K1 | G225D | - |
| MAP2K1 | G225R | - | MAP2K1 | G225S | - |
| MAP2K1 | G225V | - | MAP2K1 | T226A | - |
| MAP2K1 | T226I | - | MAP2K1 | T226K | - |
| MAP2K1 | T226P | - | MAP2K1 | T226R | - |
| MAP2K1 | R234G | - | MAP2K1 | R234T | - |
| MAP2K2 | K101E | - | MAP2K2 | K101Q | - |
| MAP2K2 | K101R | - | MAP2K2 | K101T | - |
| MAP2K2 | L119P | - | MAP2K2 | L119R | - |
| MAP2K2 | L122P | - | MAP2K2 | L122Q | - |
| MAP2K2 | L122R | - | MAP2K2 | V131E | - |
| MAP2K2 | V131G | - | MAP2K2 | I145N | - |
| MAP2K2 | I145S | - | MAP2K2 | M147K | - |
| MAP2K2 | M147R | - | MAP2K2 | R193P | - |
| MAP2K2 | R193Q | - | MAP2K2 | D194A | - |
| MAP2K2 | D194G | - | MAP2K2 | D194H | - |
| MAP2K2 | D194V | - | MAP2K2 | D194Y | - |
| MAP2K2 | D212A | - | MAP2K2 | D212G | - |
| MAP2K2 | D212H | - | MAP2K2 | D212N | - |
| MAP2K2 | D212V | - | MAP2K2 | D212Y | - |
| MAP2K2 | F213S | - | MAP2K2 | G214E | - |
| MAP2K2 | G214R | * | MAP2K2 | G214V | - |
| MAP2K2 | G214W | - | MAP2K2 | S216G | - |
| MAP2K2 | S216I | - | MAP2K2 | S216N | - |
| MAP2K2 | L219P | - | MAP2K2 | L219R | - |
| MAPK10 | L161S | - | MAPK10 | S167P | - |
| MAPK10 | P344R | - | MAPK10 | R347G | - |
| MAPK10 | R347I | - | MAPK10 | R347K | - |
| MAPK10 | R347T | - | MAPK10 | V350E | - |

| | | | | | |
|---|---|---|---|---|---|
| MAPK10 | V350G | - | MSH2 | F650S | - |
| MSH2 | I651F | - | MSH2 | I651N | - |
| MSH2 | I651S | - | MSH2 | I651T | - |
| MSH2 | N653D | - | MSH2 | N653H | - |
| MSH2 | N653I | - | MSH2 | N653S | - |
| MSH2 | N653T | - | MSH2 | N653Y | - |
| MSH2 | P670H | * | MSH2 | P670R | * |
| MSH2 | P670S | - | MSH2 | P670T | - |
| MSH2 | N671D | * | MSH2 | N671H | - |
| MSH2 | N671I | - | MSH2 | N671T | - |
| MSH2 | N671Y | * | MSH2 | M672I | - |
| MSH2 | M672K | - | MSH2 | M672R | - |
| MSH2 | M672T | - | MSH2 | G673E | - |
| MSH2 | G673R | - | MSH2 | G673V | - |
| MSH2 | G674A | * | MSH2 | G674C | - |
| MSH2 | G674D | * | MSH2 | G674R | * |
| MSH2 | G674S | * | MSH2 | G674V | - |
| MSH2 | K675E | * | MSH2 | K675I | - |
| MSH2 | K675Q | - | MSH2 | K675R | - |
| MSH2 | K675T | - | MSH2 | S676A | - |
| MSH2 | S676L | * | MSH2 | S676P | * |
| MSH2 | S676T | - | MSH2 | T677I | - |
| MSH2 | T677K | - | MSH2 | T677P | - |
| MSH2 | T677R | - | MSH2 | Y815C | - |
| MSH2 | Y815D | - | MSH2 | Y815N | - |
| MSH2 | Y815S | - | NT5C2 | K292E | - |
| NT5C2 | K292Q | - | NT5C2 | Q453E | - |
| NT5C2 | Q453K | - | NT5C2 | Q453P | - |
| PDE4D | I439F | - | PDE4D | I439N | - |
| PDE4D | I439S | - | PDE4D | I439T | - |
| PDE4D | F487S | - | PDE4D | A495E | - |
| PDE4D | A495P | - | PDE4D | S499I | - |
| PDE4D | H502D | - | PDE4D | H502L | - |
| PDE4D | H502N | - | PDE4D | H502P | - |
| PDE4D | H502R | - | PDE4D | H502Y | - |
| PDE4D | D503A | - | PDE4D | D503G | - |
| PDE4D | D503H | - | PDE4D | D503N | - |
| PDE4D | D503V | - | PDE4D | D503Y | - |
| PDE4D | H506D | - | PDE4D | H506L | - |
| PDE4D | H506N | - | PDE4D | H506P | - |
| PDE4D | H506R | - | PDE4D | H506Y | - |

| | | | | | |
|---|---|---|---|---|---|
| PDE4D | H534D | - | PDE4D | H534L | - |
| PDE4D | H534N | - | PDE4D | H534P | - |
| PDE4D | H534R | - | PDE4D | H534Y | - |
| PDE4D | H535D | - | PDE4D | H535L | - |
| PDE4D | H535N | - | PDE4D | H535P | - |
| PDE4D | H535R | - | PDE4D | H535Y | - |
| PDE4D | V538E | - | PDE4D | L600P | - |
| PDE4D | L600R | - | PDE5A | Y612C | - |
| PDE5A | Y612D | - | PDE5A | Y612H | - |
| PDE5A | Y612N | - | PDE5A | Y612S | - |
| PDE5A | L725P | - | PDE5A | L725Q | - |
| PDE5A | L725R | - | PDE5A | A779P | - |
| PDE5A | V782E | - | PDE5A | Q817E | - |
| PDE5A | Q817K | - | PDE5A | Q817P | - |
| PDE5A | Q817R | - | PKLR | Q421E | - |
| PKLR | Q421K | * | PKLR | Q421P | - |
| PKLR | Q421R | - | PKLR | L474P | - |
| PKLR | L474Q | - | PKLR | L474R | - |
| PKLR | T475A | - | PKLR | T475I | - |
| PKLR | T475N | - | PKLR | T475P | - |
| PKLR | G478A | - | PKLR | G478C | - |
| PKLR | G478D | - | PKLR | G478R | - |
| PKLR | G478S | - | PKLR | G478V | - |
| PKLR | R479C | - | PKLR | R479H | * |
| PKLR | R479L | - | PKLR | R479P | - |
| PKLR | S480A | - | PKLR | S480L | - |
| PKLR | S480P | - | PKLR | I494N | - |
| PKLR | I494S | - | PKLR | I494T | - |
| PKLR | T497A | - | PKLR | T497I | - |
| PKLR | T497N | - | PKLR | T497P | - |
| PKLR | T497S | - | PKLR | R498C | * |
| PKLR | R498G | - | PKLR | R498L | - |
| PKLR | R498P | - | PKLR | R498S | * |
| PKLR | Q501E | - | PKLR | Q501P | - |
| PKLR | W525C | - | PKLR | W525G | - |
| PKLR | W525L | - | PKLR | W525R | - |
| PKLR | W525S | - | PKLR | D528A | - |
| PKLR | D528G | - | PKLR | D528V | - |
| PKLR | D528Y | - | PKLR | V529A | - |
| PKLR | V529E | - | PKLR | V529G | - |
| PKLR | V529L | - | PKLR | D530A | - |

| | | | | | |
|---|---|---|---|---|---|
| PKLR | D530G | - | PKLR | D530H | - |
| PKLR | D530V | - | PKLR | D530Y | - |
| PKLR | R532G | - | PKLR | R532L | - |
| PKLR | R532P | - | PKLR | R532Q | * |
| PKLR | R532W | * | PKLR | V533A | - |
| PKLR | V533E | - | PKLR | V533G | - |
| PKLR | Q534P | - | PKLR | G557A | * |
| PKLR | G557C | - | PKLR | G557D | - |
| PKLR | G557R | - | PKLR | G557S | - |
| PKLR | G557V | - | PKLR | G561C | - |
| PKLR | G561R | - | PKLR | G561V | - |
| PKLR | G563A | - | PKLR | G563C | - |
| PKLR | G563D | - | PKLR | G563R | - |
| PKLR | G563V | - | PKLR | T565A | - |
| PKLR | T565I | - | PKLR | T565N | - |
| PKLR | T565P | - | PKLR | T565S | - |
| PKM | L27P | - | PKM | L27Q | - |
| PKM | L27R | - | PKM | R43G | - |
| PKM | R43L | - | PKM | R43P | - |
| PKM | R43Q | - | PKM | T45I | - |
| PKM | T45P | - | PKM | G46A | - |
| PKM | G46C | - | PKM | G46D | - |
| PKM | G46R | - | PKM | G46V | - |
| PKM | N70I | - | PKM | N70Y | - |
| PKM | R106G | - | PKM | K311E | - |
| PKM | K311M | - | PKM | K311N | - |
| PKM | K311Q | - | PKM | K311T | - |
| PKM | L353P | - | PKM | L353Q | - |
| PKM | L353R | - | PKM | D354A | - |
| PKM | D354G | - | PKM | D354H | - |
| PKM | D354N | - | PKM | D354V | - |
| PKM | D354Y | - | PKM | L431P | - |
| PKM | L431R | - | PKM | T432A | - |
| PKM | T432P | - | PKM | T432S | - |
| PKM | S434C | - | PKM | S434F | - |
| PKM | S434P | - | PKM | S434Y | - |
| PKM | G435A | - | PKM | G435C | - |
| PKM | G435D | - | PKM | G435R | - |
| PKM | G435S | - | PKM | G435V | - |
| PKM | R436G | - | PKM | R436T | - |
| PKM | S437C | - | PKM | S437F | - |

| | | |
|---|---|---|
| PKM | S437P | - |
| PKM | H464D | - |
| PKM | H464P | - |
| PKM | H464Y | - |
| PKM | G468C | - |
| PKM | G468R | - |
| PKM | G468V | - |
| PKM | I469N | - |
| PKM | I469T | - |
| PKM | P471A | - |
| PKM | P471L | - |
| PKM | P471S | - |
| PKM | W482C | - |
| PKM | W482L | - |
| PKM | W482S | - |
| PKM | R489L | - |
| PKM | R489Q | - |
| PKM | G514E | - |
| PKM | G514V | - |
| PKM | W515G | - |
| PKM | W515R | - |
| PKM | R516G | - |
| PKM | G518C | - |
| PKM | G518R | - |
| PKM | G520A | - |
| PKM | G520D | - |
| PKM | G520V | - |
| PKM | T522I | - |
| PKM | T522P | - |
| PPARG | D341G | - |
| PPARG | D341N | - |
| PPARG | D341Y | - |
| PPARG | Q342K | - |
| PPARG | Q342P | - |
| PPARG | I353N | - |
| PRPS1 | K99E | - |
| PRPS1 | K99Q | - |
| PRPS1 | K100E | - |
| PRPS1 | K100Q | - |
| PRPS1 | D101Y | - |
| PRPS1 | K102M | - |
| PKM | S437Y | - |
| PKM | H464L | - |
| PKM | H464R | - |
| PKM | G468A | - |
| PKM | G468D | - |
| PKM | G468S | - |
| PKM | I469F | - |
| PKM | I469S | - |
| PKM | F470S | - |
| PKM | P471H | - |
| PKM | P471R | - |
| PKM | P471T | - |
| PKM | W482G | - |
| PKM | W482R | - |
| PKM | R489G | - |
| PKM | R489P | - |
| PKM | G514A | - |
| PKM | G514R | - |
| PKM | W515C | - |
| PKM | W515L | - |
| PKM | W515S | - |
| PKM | G518A | - |
| PKM | G518D | - |
| PKM | G518V | - |
| PKM | G520C | - |
| PKM | G520R | - |
| PKM | T522A | - |
| PKM | T522N | - |
| PPARG | D341A | - |
| PPARG | D341H | - |
| PPARG | D341V | - |
| PPARG | Q342E | - |
| PPARG | Q342L | - |
| PPARG | Q342R | - |
| PPARG | I353S | - |
| PRPS1 | K99M | - |
| PRPS1 | K99T | - |
| PRPS1 | K100I | - |
| PRPS1 | K100T | - |
| PRPS1 | K102E | - |
| PRPS1 | K102T | - |

| | | | | | |
|---|---|---|---|---|---|
| PRPS1 | S132F | * | PRPS1 | S132Y | - |
| PRPS1 | Q135E | - | PRPS1 | Q135H | - |
| PRPS1 | Q135K | - | PRPS1 | Q135L | - |
| PRPS1 | Q135P | - | PRPS1 | Q135R | - |
| PRPS1 | N144D | - | PRPS1 | N144I | - |
| PRPS1 | N144T | - | PRPS1 | N144Y | - |
| PRPS1 | Y146D | - | PRPS1 | Y146N | - |
| PTK2 | K454E | - | PTK2 | K454Q | - |
| PTK2 | M475K | - | PTK2 | M475R | - |
| PTK2 | I483N | - | PTK2 | I483S | - |
| PTK2 | V484E | - | PTK2 | V484G | - |
| PTK2 | L486P | - | PTK2 | L486Q | - |
| PTK2 | L486R | - | PTK2 | L534H | - |
| PTK2 | L534P | - | PTK2 | L534R | - |
| PTK2 | L537P | - | PTK2 | L537Q | - |
| PTK2 | L537R | - | PTK2 | H544D | - |
| PTK2 | H544L | - | PTK2 | H544N | - |
| PTK2 | H544P | - | PTK2 | H544R | - |
| PTK2 | H544Y | - | PTK2 | R550P | - |
| PTK2 | N551D | - | PTK2 | N551H | - |
| PTK2 | N551I | - | PTK2 | N551S | - |
| PTK2 | N551T | - | PTK2 | N551Y | - |
| PTK2 | V552D | - | PTK2 | V552G | - |
| PTK2 | L562S | - | PTK2 | G563E | - |
| PTK2 | G563R | - | PTK2 | G563V | - |
| PTK2 | D564A | - | PTK2 | D564G | - |
| PTK2 | D564H | - | PTK2 | D564N | - |
| PTK2 | D564V | - | PTK2 | D564Y | - |
| PTK2 | D604A | - | PTK2 | D604G | - |
| PTK2 | D604H | - | PTK2 | D604N | - |
| PTK2 | D604V | - | PTK2 | D604Y | - |
| PTK2 | M607K | - | PTK2 | M607R | - |
| PTK2 | M607T | - | PTK2 | F608C | - |
| PTK2 | F608I | - | PTK2 | F608S | - |
| PTK2 | F608V | - | PTK2 | C611G | - |
| PTK2 | C611R | - | PTK2 | C611Y | - |
| PTPN1 | P188R | - | PTPN1 | A189P | - |
| PTPN1 | L192S | - | PTPN1 | L192W | - |
| PTPN1 | E276K | - | PYGL | L40P | - |
| PYGL | A57E | - | PYGL | A57P | - |
| PYGL | V60E | - | PYGL | L64P | - |

| | | | | | |
|---|---|---|---|---|---|
| PYGL | L64Q | - | PYGL | L64R | - |
| PYGL | W68C | - | PYGL | W68G | - |
| PYGL | W68L | - | PYGL | W68R | - |
| PYGL | W68S | - | PYGL | T71P | - |
| PYGL | T71R | - | PYGL | Q72P | - |
| PYGL | A155D | - | PYGL | A155P | - |
| PYGL | L184P | - | PYGL | L184R | - |
| PYGL | P189H | - | PYGL | P189R | - |
| PYGL | W190C | - | PYGL | W190G | - |
| PYGL | W190L | - | PYGL | W190R | - |
| PYGL | W190S | - | PYGL | E191A | - |
| PYGL | E191G | - | PYGL | E191K | - |
| PYGL | E191Q | - | PYGL | E191V | - |
| PYGL | R194C | - | PYGL | R194G | - |
| PYGL | R194P | - | PYGL | R194S | - |
| PYGL | P226L | - | PYGL | P226Q | - |
| PYGL | P226R | - | PYGL | Y227D | - |
| PYGL | Y227S | - | PYGL | T229P | - |
| PYGL | N240D | - | PYGL | N240H | - |
| PYGL | N240Y | - | PYGL | M242K | - |
| PYGL | M242R | - | PYGL | D284A | - |
| PYGL | D284G | - | PYGL | D284H | - |
| PYGL | D284V | - | PYGL | D284Y | - |
| PYGL | Q306E | - | PYGL | Q306K | - |
| PYGL | Q306P | - | PYGL | I309N | - |
| PYGL | I309S | - | PYGL | V380E | - |
| PYGL | V380G | - | PYGL | I571K | - |
| PYGL | I571R | - | PYGL | E573K | - |
| PYGL | E573Q | - | PYGL | A610D | - |
| PYGL | A610G | - | PYGL | A610P | - |
| PYGL | A610V | - | PYGL | D770A | - |
| PYGL | D770G | - | PYGL | D770H | - |
| PYGL | D770N | - | PYGL | D770V | - |
| PYGL | D770Y | - | PYGM | T71P | - |
| PYGM | T71R | - | PYGM | I309N | - |
| PYGM | I309S | - | PYGM | A610D | - |
| PYGM | A610G | - | PYGM | A610P | - |
| PYGM | A610V | - | RRM1 | R6G | - |
| RRM1 | R6L | - | RRM1 | R6P | - |
| RRM1 | R6Q | - | RRM1 | L56P | - |
| RRM1 | L56Q | - | RRM1 | K88E | - |

| | | | | | |
|---|---|---|---|---|---|
| RRM1 | D226A | - | RRM1 | D226G | - |
| RRM1 | D226V | - | RRM1 | S227I | - |
| RRM1 | I228N | - | RRM1 | I231S | - |
| RRM1 | K243E | - | RRM1 | I255S | - |
| RRM1 | R256G | - | RRM1 | R256L | - |
| RRM1 | R256P | - | RRM1 | R256Q | - |
| RRM1 | I262S | - | RRM1 | G264E | - |
| RRM1 | G264R | - | RRM1 | S269F | - |
| RRM1 | S269P | - | RRM1 | S269Y | - |
| RRM1 | V286E | - | RRM1 | Q288E | - |
| RRM1 | Q288K | - | SAMHD1 | D120A | - |
| SAMHD1 | D120G | - | SAMHD1 | D120H | - |
| SAMHD1 | D120V | - | SAMHD1 | D120Y | - |
| SAMHD1 | I136N | - | SAMHD1 | I136S | - |
| SAMHD1 | Q142E | - | SAMHD1 | Q142K | - |
| SAMHD1 | Q142P | - | SAMHD1 | R145P | * |
| SAMHD1 | V156D | - | SAMHD1 | L169P | - |
| SAMHD1 | L169Q | - | SAMHD1 | L169R | - |
| SAMHD1 | R333G | - | SAMHD1 | R333P | - |
| SAMHD1 | R333S | - | SAMHD1 | K354E | - |
| SAMHD1 | H376D | - | SAMHD1 | H376L | - |
| SAMHD1 | H376N | - | SAMHD1 | H376P | - |
| SAMHD1 | H376R | - | SAMHD1 | H376Y | - |
| SAMHD1 | R451G | - | SAMHD1 | R451S | - |
| SERPINC1 | P112A | - | SERPINC1 | P112H | - |
| SERPINC1 | P112L | - | SERPINC1 | P112R | - |
| SERPINC1 | P112S | * | SERPINC1 | P112T | * |
| SERPINC1 | L113P | - | SERPINC1 | L113R | - |
| SERPINC1 | S114I | - | SERPINC1 | S114N | * |
| SERPINC1 | M121K | * | SERPINC1 | M121R | - |
| SERPINC1 | M121T | - | SERPINC1 | G125A | - |
| SERPINC1 | G125C | - | SERPINC1 | G125D | * |
| SERPINC1 | G125R | - | SERPINC1 | G125S | - |
| SERPINC1 | G125V | - | SERPINE1 | L66S | - |
| SERPINE1 | V122D | - | SERPINE1 | Q227P | - |
| SIRT3 | Q228E | - | SIRT3 | Q228K | - |
| SIRT3 | Q228P | - | SIRT3 | D231A | - |
| SIRT3 | D231G | - | TTR | P31H | - |
| TTR | P31L | - | TTR | P31R | - |
| TTR | L32P | * | TTR | L32Q | - |
| TTR | L32R | - | TTR | M33I | - |

| | | | | | |
|---|---|---|---|---|---|
| TTR | M33K | - | TTR | M33R | - |
| TTR | P44A | - | TTR | P44H | - |
| TTR | P44L | * | TTR | P44R | - |
| TTR | P44S | * | TTR | P44T | - |
| TTR | A45D | - | TTR | A45G | - |
| TTR | A45P | - | TTR | A45S | - |
| TTR | A45T | * | TTR | A45V | - |
| TTR | V52A | * | TTR | V52E | - |
| TTR | V52G | - | TTR | L78H | * |
| TTR | L78P | - | TTR | L78R | * |
| TTR | F84C | - | TTR | F84I | - |
| TTR | F84S | * | TTR | F84V | - |
| TTR | H108D | - | TTR | H108L | - |
| TTR | H108N | - | TTR | H108P | - |
| TTR | H108R | * | TTR | E109G | - |
| TTR | E109V | * | TTR | F115C | - |
| TTR | F115I | - | TTR | F115S | - |
| TTR | F115V | - | TTR | F115Y | - |
| TTR | A117D | - | TTR | A117G | * |
| TTR | A117P | - | TTR | N118I | - |
| TYMS | Q160E | - | TYMS | Q160K | - |
| TYMS | Q160L | - | TYMS | Q160P | - |
| TYMS | Q160R | - | TYMS | I178N | - |
| TYMS | I178S | - | TYMS | M179K | - |
| TYMS | M179R | - | TYMS | W182C | - |
| TYMS | W182G | - | TYMS | W182L | - |
| TYMS | W182R | - | TYMS | W182S | - |
| TYMS | N183D | - | TYMS | N183I | - |
| TYMS | N183T | - | TYMS | N183Y | - |
| TYMS | L192P | - | TYMS | L192Q | - |
| TYMS | L192R | - | TYMS | H196D | - |
| TYMS | H196L | - | TYMS | H196R | - |
| TYMS | R215G | - | TYMS | R215I | - |
| TYMS | R215K | - | TYMS | S216L | - |
| TYMS | S216P | - | TYMS | S216T | - |
| TYMS | S216W | - | TYMS | Y258C | - |
| TYMS | Y258D | - | TYMS | Y258H | - |
| TYMS | Y258N | - | TYMS | Y258S | - |
| UGDH | T131A | - | UGDH | T131I | - |
| UGDH | T131K | - | UGDH | T131P | - |
| UGDH | T131R | - | UGDH | E161A | - |

| | | | | | |
|---|---|---|---|---|---|
| UGDH | E161G | - | UGDH | E161K | - |
| UGDH | E161Q | - | UGDH | E161V | - |
| UGDH | F162C | - | UGDH | F162I | - |
| UGDH | F162S | - | UGDH | F162V | - |
| UGDH | L163P | - | UGDH | L163Q | - |
| UGDH | L163R | - | UGDH | E165G | - |
| UGDH | E165K | - | UGDH | E165Q | - |
| UGDH | E165V | - | UGDH | K220E | - |
| UGDH | K220I | - | UGDH | K220Q | - |
| UGDH | K220R | - | UGDH | K220T | - |
| UGDH | N224D | - | UGDH | N224H | - |
| UGDH | N224I | - | UGDH | N224S | - |
| UGDH | N224T | - | UGDH | N224Y | - |
| UGDH | L227F | - | UGDH | L227H | - |
| UGDH | L227P | - | UGDH | L227R | - |
| UGDH | I231K | - | UGDH | I231R | - |
| UGDH | I231T | - | UGDH | R260G | - |
| UGDH | R260I | - | UGDH | R260K | - |
| UGDH | R260S | - | UGDH | R260T | - |
| UGDH | F265I | - | UGDH | F265S | - |
| UGDH | F265V | - | UGDH | L266P | - |
| UGDH | L266Q | - | UGDH | L266R | - |
| UGDH | S269C | - | UGDH | S269I | - |
| UGDH | S269N | - | UGDH | S269T | - |
| UGDH | F272S | - | UGDH | G273A | - |
| UGDH | G273C | - | UGDH | G273D | - |
| UGDH | G273R | - | UGDH | G273S | - |
| UGDH | G273V | - | UGDH | C276F | - |
| UGDH | C276G | - | UGDH | C276R | - |
| UGDH | C276S | - | UGDH | C276Y | - |
| UGDH | F277C | - | UGDH | F277S | - |
| UGDH | F277V | - | UGDH | F338I | - |
| UGDH | F338S | - | UGDH | F338V | - |
| UGDH | K339E | - | UGDH | K339I | - |
| UGDH | K339Q | - | UGDH | K339T | - |

* is the pathogenic mutation recorded by Clivar;

– is the highly potential pathogenic mutation predicted by MetaMutPre.
